# Optimizing non-invasive assessment of gastric activity: a fully automated channel-wise pipeline for multi-electrode electrogastrography

**DOI:** 10.64898/2026.10.28.762445

**Authors:** Iannone Alessandro, Panasiti Maria Serena, Aglioti Salvatore Maria, Della Penna Stefania

**Affiliations:** Department of Psychology, Sapienza University of Rome, Rome, Italy; IRCCS, Santa Lucia Foundation, Rome, Italy; Sapienza University of Rome and CLN2S, Fondazione Istituto Italiano di Tecnologia (IIT), Rome, Italy; Department of Neuroscience, Imaging, and Clinical Sciences, University “G. d’Annunzio” of Chieti-Pescara, Chieti, Italy; Institute for Advanced Biomedical Technologies, University “G. d’Annunzio” of Chieti-Pescara, Chieti, Italy; UdA-TechLab, University “G. d’Annunzio” of Chieti-Pescara, Chieti, Italy

**Author notes:** These authors contributed equally to this work (Aglioti Salvatore Maria, Della Penna Stefania). Corrisponding author.

**Keywords:** Electrogastrography, channel-wise preprocessing, artifact rejection, data driven analysis, gastric fingerprint

## Abstract

Electrogastrography (EGG) is a non-invasive, and affordable technique for investigating gastric physiology. However, its broader adoption is hindered by low signal-to- noise ratios and the lack of standardized, objective procedures for channel-wise artifact rejection and signal quality assessment. To address these limitations, we developed a fully automated, data-driven pipeline (Gastric Signal Processing Toolbox – GSPT) for channel-wise EGG preprocessing and analysis, designed to preserve physiologically-relevant information even under high-noise conditions.

GSPT exploits high-resolution time-frequency representations, to automatically identify and remove motion-related and electronic artifacts through entropy-based analysis of the signal’s instantaneous energy and robust outlier detection through integration of multiple anomaly-detection strategies. The resulting signals are characterized through a channel-wise gastric fingerprint, comprising established EGG measures, such as dominant frequency, together with novel descriptors such as Renyi entropy. Channel quality relays on Monte Carlo surrogate analysis, while recording-level quality is evaluated through dominant frequency consistency across channels.

GSPT further provides ridge-based estimates of instantaneous amplitude and phase and quantifies inter-channel interactions through Total Correlation, enabling the construction of a global gastric fingerprint.

Validation on datasets containing controlled motion artifacts demonstrated robust signal recovery with improved signal-to-noise ratios at the dominant frequency. Artifact identification, assessed using precision and recall, was comparable or better than existing approaches. Gastric fingerprints showed high stability within subjects and across recording sessions. As a freely available, one-click MATLAB toolbox, GSPT reduces manual bias and standardizes EGG preprocessing and analysis, even in low-density recordings, potentially providing a reproducible framework for investigating gastric functions in psychological, neurological and pathological conditions.

## I. Introduction

Electrogastrography (EGG) is a non-invasive technique used to monitor gastric myoelectrical activity (GMA) via cutaneous electrodes. Since its inception in the early 1920s, EGG has been used to record the gastric slow wave—a rhythmic electrical potential generated by the Interstitial Cells of Cajal (ICC) at approximately 0.05 Hz—as well as the activity associated with gastric peristaltic contractions.

EGG has attracted growing interest of the scientific community as a valuable tool for investigating gastric activity because of its high sensitivity to a wide range of physiological and pathological conditions and of its relatively simple, cheap and non-invasive setup usually comprising only a few surface electrodes. EEG has been proposed for probing conditions such as gastroparesis, Parkinson’s disease, and diabetes mellitus [1, 2, 3], as well as for research in psychology and neuroscience [4, 5], including the study of complex brain–gut interaction networks [6, 7].

Despite its potential, EGG’s clinical reliability is limited by several challenges. First, gastric signals occur at ultra-low frequencies and are highly susceptible to electronic interference and motion artifacts. Second, participants’ movements often produce spectral components that directly overlap with the EGG bands, making simple filtering insufficient. Furthermore, EGG is typically recorded with few channels (1–4), precluding advanced spatial filtering or high-density decomposition like ICA or similar that are usually applied on larger channel numbers for artifact rejection. A standard approach for processing EGG signals relies on static power spectra or inter-channel dominant frequency coherence methods [4]. However, this approach is suboptimal, since it does not take into consideration the temporal dynamics of the gastric signal and is strongly operator-dependent. Alternative methods, such as decomposition methods (e.g., DWT, EMD, NA-MEMD) [8] considerably improve the information provided on EGG dynamics, but they may not be well suited to handling electronic and motion artifacts, given their sensitivity to spikes and abrupt transients [9]. An alternative method based on LMMSE estimators [10], has been proposed and adopted in several studies. This approach is advantageous for avoiding the need to discard artifact-contaminated data segments and has been integrated in a commercial low-noise EGG multichannel system. It relies on assumptions of local stationarity to identify non-stationary artifacts and considerably reduce them on the raw signal.

As an alternative, complete approach for EGG analysis, here we propose a pipeline, namely GSPT (Gastric Signal Processing Toolbox), that provides a comprehensive, fully automated framework for systematic artifact/noise removal, quality assessment and feature extraction at the single-channel level. GSPT relies on Superlet transform - a recently proposed technique for generating high-resolution time-frequency representations (TFRs) of signals [11] – to obtain a high-resolution TFR even at the ultra-low gastric frequencies. From the TFR, the dynamics of the instantaneous energy is analysed through entropy and dispersion measures (e.g. kurtosis) to identify artifactual channels. To offer flexibility, this preliminary step is optional, allowing users to adopt a less conservative preprocessing approach if desired. Contaminated segments are then removed by applying advanced time series anomaly detection (TSAD) to preserve the TFR topography in the artifact-free intervals and obtain information even in low SNR recordings.

The efficacy of the pipeline is tested on real EGG recordings with controlled movement artifacts using modern TSAD evaluation metrics for a quantification of algorithmic performance. Furthermore, the same data and metrics are used to compare the performance of GSPT against a standard Hampel filter and the LMMSE approach. An additional comparison is conducted based on the algorithms’ ability to disentangle the power in the dominant versus side bands.

Moreover, unlike conventional approaches that rely on selecting a single “most representative” channel per measurement—a practice that risks sacrificing information and statistical power—our method operates channel-wise and accommodates an arbitrary number of channels, ensuring comprehensive data utilization.

The pipeline provides classical EGG metrics (e.g., dominant frequency, fraction of normogastria) from the low-noise TFR, tests their significance via a Monte Carlo surrogate test and dominant frequency consistency between channels, and also estimates quantitative descriptors not currently considered in the standard EGG literature (e.g., spectral entropy, Renyi entropy, active information storage) that may be useful to provide a more complete characterization of gastric activity. Notably, the pipeline allows the user to select a subset of features or even add further ones. These features collectively enable the construction of a fingerprint for each EGG channel, that might allow capturing unique physiological signatures that would remain elusive to traditional analysis. Moreover, the pipeline extracts instantaneous gastric amplitude, frequency, and phase for each channel.

Lastly, in multi-channel recordings, the pipeline can calculate multivariate and nonlinear correlation metrics across all channels via Total Correlation and multivariate entropy, providing insights into the spatial coordination and dynamical interdependencies of gastric electrical activity.

## II. METHODS

The pipeline presented in this work is designed for the analysis of individual channels, making it particularly suited to scenarios with a limited number of EGG channels, where methods like ICA can’t be used. Nevertheless, GSPT can also be extended to multichannel recordings. The general workflow of GSPT is illustrated in Fig 1. The pipeline begins by removing excessive spikes, detrending and filtering the EGG data in a selected frequency band including the gastric activity (0.016-0.16 Hz), for each channel. Then, we apply the Fractional Adaptive Superlet Transform (FASLT), a modified version of the Superlet Transform (SLT), offering superior time-frequency resolution compared to traditional TFR methods, a critical feature for the pipeline’s effectiveness in analysing the ultra-slow frequencies of the EGG data.

**Figure 1:**
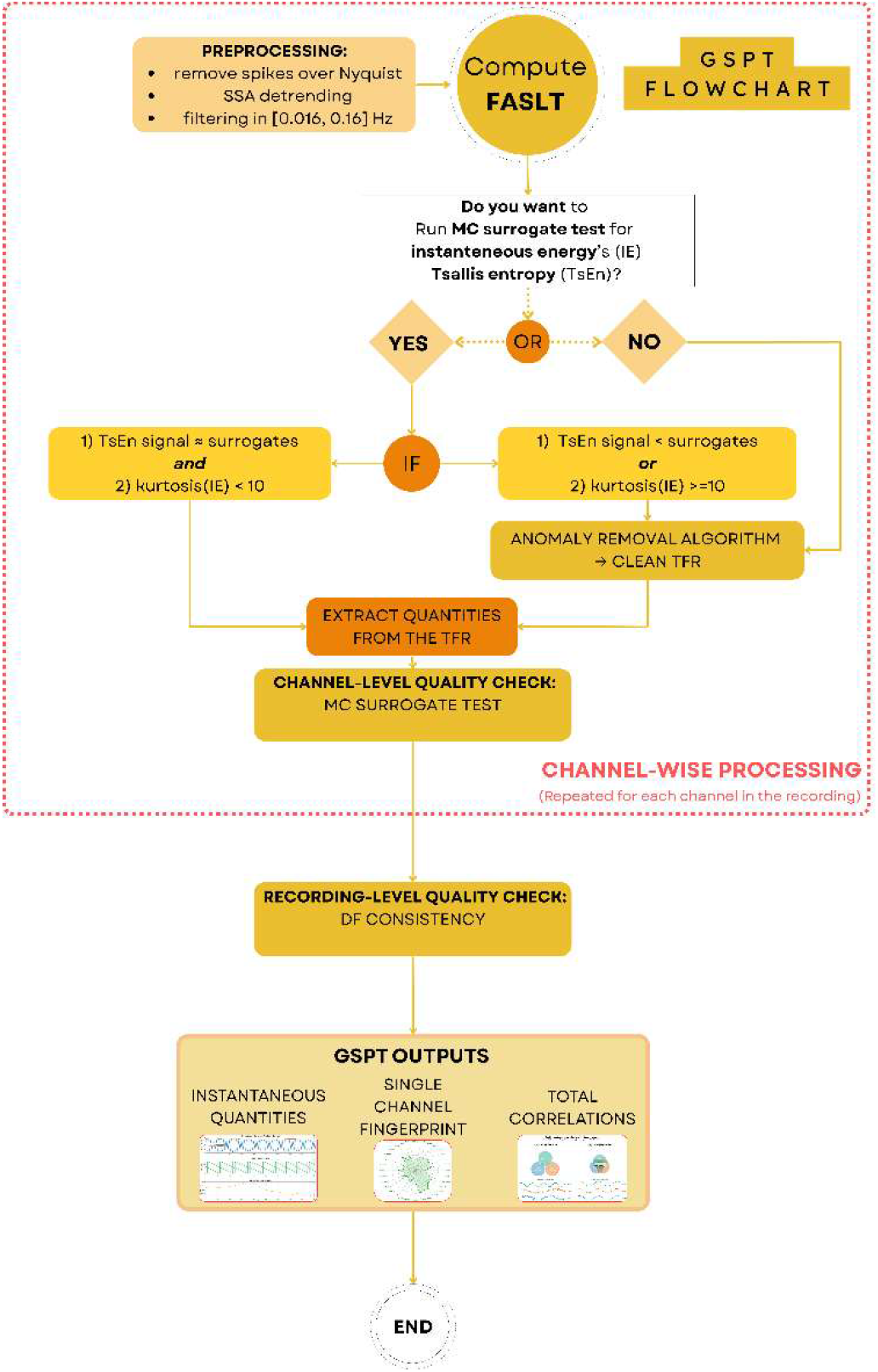
Overview of the GSPT workflow for channel-wise EGG processing and recording-level analysis.

From the FASLT-derived TFR, we then calculate a curve called Instantaneous Energy (IE), its kurtosis and its Tsallis entropy (TE). These metrics enable the detection of time-localized artifacts in the EGG recording, determining whether artifact removal algorithms need to be applied. Notably, GSPT offers users the flexibility to either run this preliminary check on the IE - resulting in a more conservative analysis pipeline - or bypass it directly to the anomaly removal algorithm for a less conservative version that identifies more anomalies but increases the risk of false positives. Conversely, the conservative approach yields fewer anomalies, isolating only those that impact the signal most severely, but carries a higher risk of false negatives. The selection between these alternatives is left to the user’s discretion, depending on the specific requirements of their analysis. After the conditional processing with the artifact removal algorithm, we can obtain an artifact free TFR. Eventually, we can extract different channel features building a channel fingerprint that characterize many aspects of the recording. The quality of the signal underlying these features is assessed through Monte Carlo statistical test at the channel-level and via the consistency of the dominant frequency across the channels at the entire run level. As additional outputs, instantaneous phase and amplitude, together with measures of channel interactions are provided.

### A. PREPROCESSING STEPS

The first step is to eliminate spikes and/or jumps whose frequency content exceeds the Nyquist frequency, that might occur in the data due to problems in the readout electronics or in the acquisition setup. This approach prevents aliasing and distortion in the TFR, that might affect even the slow frequencies of the gastric signal. First, we evaluate whether the amplitude of the spike/jump, defined as the absolute difference between consecutive signal samples, implies a violation of the Nynquist’s teorem (mathematical details can be found in the Supplementary materials - Section I: Spectral content of jumps). In this case, the spike/jump is flagged as an artifact and is corrected by subtracting the mean of the preceding f_s_/10 samples (f_s_ being the original sampling frequency). In the second step, the whole signal is then downsampled to 10 Hz, discarding the first and last four samples to eliminate filtering-related edge effects.

Signals are then detrended via Singular Spectrum Analysis (SSA) [12] using a 60-second window to preserve bradygastric components while adaptively removing low-frequency noise. A zero-phase, band-pass Chebyshev Type II cascade (0.014–0.18 Hz; orders 6 and 10) is then applied.

Finally, the TFR is computed via FASLT over the 0.016–0.16 Hz range (0.001 Hz frequency bin). We use superlets with an initial wavelet cycle of 3 and a super resolution order interval of 3–5.

### B. EVALUATION OF CHANNEL CONTAMINATION

In the more conservative approach, evaluation of potential channel contamination by transient artifacts is based on the analysis of the IE in the frequency band of interest, estimated from the TFR through the following formula:

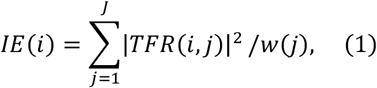

where *TFR(i, j)* is the signal TFR matrix with dimensions time X frequency, obtained by FALST, *w* is the vector of frequencies, *i* and j are the time and frequency indices, respectively. Superlets are L1 normalized instead of L2, to preserve amplitude, hence the factor 1/w(j) in the formula (see Supplementary Materials in [11]).

If there are time localized artifacts in the signal, these produce peaks in the IE. To evaluate the potential contamination of the specific channel, we first analyze the minimum TE, signalling if artifacts affect the signal and we need to apply the anomaly detection algorithm. The minimum TE is defined as:

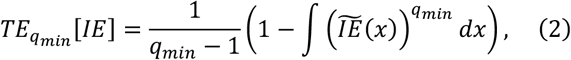

Where 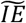 is the normalized IE curve, *q*_min_ is an adaptive parameter, selected for each channel and recording to minimize TE. While flat IE curves yield high entropy, the energy and temporal concentration of movement artifacts result in low TE. To identify noisy channels, instead of using a fixed threshold, that might be unsatisfying due to high inter-subject variability and the subjectivity of empirical tuning, we utilize Monte Carlo surrogate testing [13]. Specifically, we label recordings as affected by artifacts through comparison of the observed TE against a data-driven null distribution obtained via random permutation surrogates. Segments indistinguishable from surrogates undergo a secondary safety check using a conservative kurtosis threshold (> 10). Additional details are provided in the Supplementary materials, Section II: Tsallis Entropy.

### C. LOCALIZING AND REJECTIG ANOMALIES

Once a channel is labelled as bad in the more conservative approach, or always in the less conservative one, we pinpoint anomalous intervals using a sequence of three complementary algorithms on EGG time series, in the following sequence: Tukey Fences, Connectivity Outlier Factor (COF) [14], and Robust Random Cut Forest (RRCF) [15].

1. Tukey Fences (Local & Global): A local Tukey method (100s sliding window, 99.9% overlap, 1.5 Inter Quartile Range threshold) detects dynamic outliers (see Supplementary Materials, Section III, for an example) but can yield false negatives at signal edges and false positives during physiological amplitude shifts. To mitigate this risk, we also apply a global Tukey to the entire recording.
2. COF Validation: To handle the non-stationarity that biases global thresholds, global Tukey flags are retained only if confirmed by COF.
3. RRCF Refinement: Finally, an RRCF model is deployed to minimize false positives. We objectively set its contamination fraction based on the proportion of points previously flagged by the union of the results obtained by local Tukey and the [Global Tukey ∩ COF]. To account for the stochastic nature of RRCF, which can cause minor shifts in the boundaries of the detected anomalies (typically within one or a few samples), we set the RRCF seed using the Mersenne Twister random generator with default seed 0. This guarantees reproducibility across runs.

The final set of anomalies consists of epochs detected by the RRCF that also appear in the preliminary detection set. This multi-method integration aligns with recent trends in robust time-series analysis [16], ensuring a conservative, data-driven artifact removal that preserves genuine gastric dynamics even at recording edges.

Following temporal localization, anomalies are managed via a reconstruct-then-mask strategy designed to minimize spectral distortion and edge effects in the TFR, that might be induced by simply removing the artifact epochs from the signal and are particularly detrimental to ultra-low frequency EGG analysis. This strategy follows two stages:

1. AR Gap-Filling: Missing intervals are reconstructed using autoregressive (AR) model estimates (via the MATLAB’s function fillgaps). This procedure blends forward and backward extrapolations using a linear weighted mean. To account for EGG non-stationarity, AR models are estimated using a 90-second window adjacent to each gap, instead of the whole recording. The model order is selected automatically by minimizing the Akaike Information Criterion (AIC).
2. TFR Masking: A new FASLT-based TFR is computed from the reconstructed signal. Finally, the TFR segments corresponding to the original anomalous intervals are removed.

### D. GSPT VALIDATION

To ensure a rigorous TSAD benchmarking based on ground-truth labeled anomalies, we applied a controlled experimental protocol where participants moved only upon explicit request and remained still otherwise during a continuous EGG recording, thereby enabling the collection of anomaly timestamps. We recorded EGG from 9 participants instructed to move only upon explicit request allowing for timestamping of 248 movement events (mean duration: 4.03 s, sd: 1.58 s) with a total of 17 runs.

Quantitatively assessing TSAD performance is non-trivial due to the inherent imprecision in labeling and the temporal smearing introduced by gastric-range filtering. To accommodate these uncertainties, we adopted segment-wise metrics (Precision and Recall) [17]. In this framework, a predicted anomaly is classified as a True Positive if it overlaps with any portion of a ground-truth segment.

We thus compared precision and recall of the conservative and non-conservative approaches with a traditional Hampel filter (with a standard threshold of 3 MAD) and with LMMSE. However, a challenge remains: movement artifacts frequently consist of high-frequency components that do not significantly impact the gastric frequency band (0.016–0.16 Hz).

Consequently, using all the timestamped events can be misleading, biasing the false negative rate. To mitigate this bias, we employed the LMMSE-like estimator [10] to label the events as false or true ones through a quantitative assessment of their impact on the gastric band. Specifically, we first applied FALST for both the raw and the LMMSE-processed signals, normalized these TFRs by the square root of their respective dominant powers and computed the difference between the absolute values of the normalized raw and LMMSE TFRs. To identify time-frequency epochs of true artifacts in the gastric frequency band, this difference was compared against TFRs generated from 100 randomly shuffled surrogates of the original signal, which were normalized using the square root of the raw signal’s dominant power. Time intervals among the timestamped ones were flagged as true artifacts epochs if the TFR difference at any point exceeded the surrogate TFR mean plus two standard deviations. If a labeled artifact was not identified by this threshold, it was classified as a false negative, but it indicated that the artifact’s impact on the gastric frequency range was negligible relative to the noise floor. Hence, we moved the false negatives of each other approach (conservative GSPT, non-conservative GSPT, Hampel) that overlapped with the LMMSE false negatives to true negative events. As a consequence, comparison with LMMSE was based on precision only.

As an additional validation measure, we evaluated the impact of GSPT on the dominant power, a critical quantity in the EGG analysis. Specifically, we estimated the ratio of dominant power (DF±0.015 Hz) to background power-defined here as the mean power in the frequency band [DF + 0.015 Hz, 0.16 Hz], on the same dataset containing controlled artifacts. A higher value for this quantity indicates better peak distinction from unwanted signals, which are primarily related to movement artifacts that tend to produce a flatter power spectral density. We compared this quantity across the signal before cleaning, after GSPT, after Hampel filter, and after LMMSE cleaning. We selected the DF on the GSPT-cleaned data, and used the very same frequency for the other conditions.

### E. FEATURE EXTRACTION: EGG FINGERPRINT

Once a clean TFR is obtained, GSPT evaluates a set of features forming a multidimensional fingerprint to characterize the selected EGG channel. These features comprise classical spectral metrics—such as dominant frequency (DF), power, and the fraction of normogastria— and other features novel for the EGG field (see Table S1, Section IV and Table S2, Section V in the Supplementary Materials). For the user, it is possible to select different sets of features among the ones proposed or add other features to form the individual fingerprint, based on the specifical research questions. Concerning the first set of metrics, unlike traditional methods that rely on arbitrary binning of the frequency dimension, we estimate the percentage of normogastria (0.033–0.067 Hz) directly from Instantaneous Frequency (IF) curves. These ridges are extracted and delineated on the Superlet TFR using a global optimization approach adapted from the Synchrosqueezing Toolbox [18, 19]. In addition to relative power across standard frequency bands, we introduce adaptive band partitioning centered on the individual DF (Table S1, Section IV, Supplementary materials).

The novel features can be grouped in 4 categories (Table S2, Section V, Supplementary materials):

- Measures of temporal and spectral dynamics: Total gastric energy and its temporal variability, spectral centroid, spread, spectral entropy and Tsallis entropy for the power spectrum density.
- 1D entropy measures: Dispersion, and Sample entropy for the raw signal in the gastric band and its instantaneous energy.
- 2D TFR entropies: Renyi and Gradient entropies, Hoyer measure.
- Non-linear dynamics: Correlation dimension, Lyapunov exponent, Active Information Storage (AIS), and Detrended Fluctuation Analysis (DFA) exponent. [20]

These measures could be used to characterize system dynamics, and in our case could be useful for capturing the complex, non-linear dynamics often missed by standard spectral analysis, and might support the understanding gastric dysrhythmias associated with psychological or pathological states. In this view, we tested the consistency of the EGG fingerprint. We recruited 61 healthy participants (normal BMI, fasted ≥ 3 h, no GI symptoms), who provided two 15-minute resting-state EGG recordings each on separate days (Sessions S_1_ and S_2_). Data were acquired via a bipolar montage following established protocols [4], with the expectation of stable normogastric activity. We examined the consistency of the full EGG fingerprint at three distinct levels:

- Inter-channel: measures the internal consistency of an individual’s gastric signal across the sensor array.
- Intra-subject: evaluates temporal stability by correlating the same channels for each participant across different Sessions (S_1_ Vs S_2_).
- Inter-subject: assesses the similarity between different participants’ fingerprints (mean correlation between a subject and all others per channel).

Analyses were performed using both the full 32-feature fingerprint and a minimal three-feature fingerprint comprising DF, dominant power, and fraction of normogastria, selected as conventional measures commonly reported in EGG research. In addition to single channel scalar metrics, GSPT provides also estimates of single channel instantaneous quantities, such as phase, frequency, and amplitude of the fundamental gastric mode that could also be used to investigate the coupling between EGG and signals generated by other organs, e.g. brain [6]. GSPT derives these quantities directly from the wavelet TFR using ridge extraction and signal reconstruction [21], limiting the analysis in the frequency domain to a multiplicative window around the DF to increase precision (DF/1.5 – DF*1.5). Moreover, these instantaneous quantities, alongside filtered EGG data in the whole gastric frequency band, instantaneous energy, instantaneous spectral entropy (normalized Shannon entropy of the instantaneous Superlet power distribution) and ridge points extracted from the Superlet TFR, form the basis for our inter-channel coherence analysis. We quantified global synchronization across the montage using Multivariate Permutation and Dispersion entropies [22] and Total Correlation (TC), the multivariate generalization of mutual information that expresses, as a single scalar, the total statistical dependence, both linear and non-linear, shared among all channels [23]. For channels X_1_,…,X_n_, TC is defined as the difference between the sum of the marginal entropies and the joint entropy, TC = Σi S(Xi) − S(X_1_,…,X_n_), and vanishes if and only if the channels are jointly independent. We estimated the entropy terms with the matrix-based Rényi α-order entropy functional (α = 2), which derives all quantities from Gaussian-kernel Gram matrices without explicit density estimation and captures non-linear dependence through the kernel [24, 25].

While TC is inherently sensitive to inter-electrode distance, the adoption of a standardized montage across all participants limits this geometric bias. Under this consistent configuration, high TC indicates a redundant, coherent system driven by a stable pacemaker. Conversely, a significant reduction in TC signals the physiological decoupling and spatial heterogeneity characteristic of dysrhythmic states [26], or transient disruptions induced by emotional and psychological stress.

Finally, we complement TC with multivariate entropy to differentiate between highly synchronized, simple dynamics and complex, unpredictable joint behaviors. Similarly to the single-channel fingerprint analysis, we also quantitatively compared the intra-subject and inter-subject total correlation fingerprints (comprising total correlations and multivariate entropies for each subject’s 4-channel recording) for the same group of 61 participants

### F. QUALITY-CHECK VIA SURROGATE TEST AND DOMINANT FREQUENCY CONSISTENCY

GSPT includes a quality-check framework to validate the spectral quantities obtained from EGG recordings. Since EGG signals often have a low signal-to-noise ratio, noise can produce spectral patterns that mimic physiological gastric activity and might lead to false-positive results. Although DF is generally more robust than dominant power, it can still be distorted by noise. For instance, pure electronic noise can exhibit a ‘normogastric’ peak that can be distinguished from genuine physiological activity only with quantitative validation. (see Fig S3a and Fig S3b, Section VI in Supplementary materials). To address this issue, GSPT applies a Monte Carlo (MC) surrogate test [13] using a MATLAB version of the wavelet-based surrogates (IAAWFT) based on the implementation of the *dualtrees* R package [27] with default parameters, which account for potential signal non-stationarity. Experimental and surrogate power spectrum densities are compared using two metrics—skewness and Hoyer measure (details in Section VII – Spectral skewness and Hoyer measure in Supplementary Materials)—that are independent from the exact DF value and adapt to possible shifts from normogastria. These two quantities detect if the PSD is concentrated around a few peaks (typical of gastric activity) or spread across multiple peaks (typical of noise).

Signals are considered significantly different from noise if the Simes-adjusted empirical p-value [28] is below 0.05. GSPT also performs whole-recording quality-check based on the spatial consistency of DF across EGG channels. Since healthy gastric rhythms typically exhibit consistent DF values across electrode locations, GSPT automatically assesses recording quality by computing a corrected maximum difference (or range) Δ_max_ between DF values across channels and comparing it with a threshold, that is selected by the user. In this work, we explore three candidate thresholds to show possible outcomes of the analysis of DF consistency: 0.005 Hz, 0.01 Hz and 0.02 Hz. Recordings can be classified as either reliable (their Δ_max_ is below the threshold) or corrupted (Δ_max_ above the threshold). Assuming the measured DF in high-quality recordings represents a single underlying physiological oscillator subjected to additive Gaussian noise, we can mathematically relate the experimental Δ_max_ to the sample standard deviation *s* via the following equation:

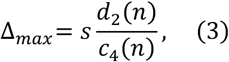

where *n* represents the number of recording channels, *c*_4_ is the Shewhart constant yielding an unbiased estimate of the population standard deviation from *s*, and *d*_2_ is the Tippett constant, which relates the expected range of a normally distributed sample to its standard deviation [29].To account for recordings in which only some channels are noisy, GSPT employs a recursive channel-selection procedure that iteratively removes channels contributing most to DF Δ_max_ until the latter falls below the threshold, or a minimum channel “quorum” is reached (at least 50% of the initial channels and no fewer than three). If the range of the remaining channels is below the threshold, they are considered to contain reliable physiological information.

The user can choose to use either MC or SD consistency alone, or to combine both in a sequence to obtain a more stringent selection criterion.

## III. RESULTS

### A. PREPROCESSING

An example of a jump on the recorded signal and its correction is shown in Fig 2. To quantify the impact of this correction, we compared these two signals, which remain identical up to the point of the jump. Over the first 900 seconds of the recording, the sample-to-sample difference (at a 1000 Hz sampling rate) is 1.36 ± 1.76 µV. In the raw signal, the samples immediately before and after the jump differ by 880.39 µV, whereas after the correction, this difference becomes 0.26 µV, that is compatible with the regular inter-sample difference (see insert in Fig 2).

**Figure 2:**
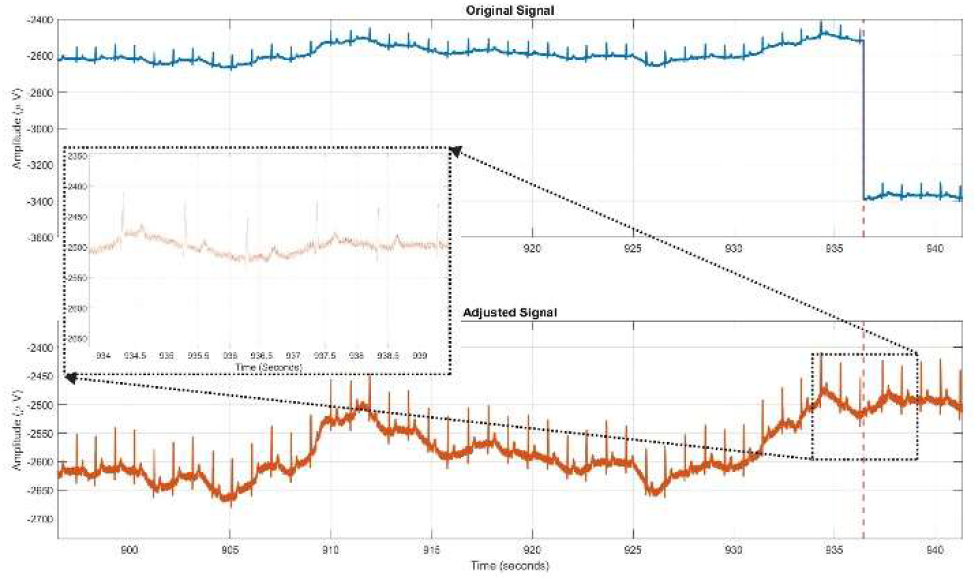
A real recorded EGG signal with a jump around 935 seconds, probably due to electronic problems. The jump is correctly identified and corrected.

### B. TSALLIS ENTROPY ON INSTANTANEOUS ENERGY

To evaluate the performances of Tsallis Entropy on IE, we compared our dataset (with controlled artifacts) with a group consisting of recordings on a subset of the same group (6 runs) and recordings from a different group (6 runs), all with participants at rest (Fig S4, Section VIII, A, Supplementary materials). We run an ART random effects model to account for the correlation between the EGG channels of the same participant and for non-normality of residuals. Although GSPT applies the “bad channel” label based on MC comparison, this analysis confirmed that Tsallis Entropy is promising for labeling noisy channels, since recording with movement artifacts showed a significantly lower Tsallis entropy (p= 0.036), that is compatible with the occurrence of movement-induced local energetic spikes.

### C. EFFICACY OF THE ARTIFACT REJECTION ALGORITHM

To provide an example of GSPT performances in rejecting artifact, Fig 3 depicts the TFR of a representative EGG signal with a visible artifact before and after GSPT application. In the upper part of Fig. 3, the raw signal shows a clear artifact at about 530 s and a less distinguishable one at about 400 s. These artifacts, especially the larger one, introduce excess power in the TFR down to the ultra-low frequency band, as shown in Fig.3, middle panel. This significantly biases the DF, which shifts to 0.0206 Hz, while only a minor secondary peak is visible in the PSD at approximately 0.04 Hz. The Tsallis entropy of the raw data IE is about 0.71, significantly lower than the surrogate data (p<0.004). Following GSPT artifact rejection (bottom panel), however, the IE entropy increases to 0.99 and a dominant frequency at precisely 0.0473 Hz emerges, while the ultra-low frequency contribution decreases. By calculating the ratio between the PSD at 0.0473 Hz and the PSD at 0.0206 Hz, we observe an increase from 0.50 to 1.24 after the GSPT correction. Additional examples can be found in Supplementary Material, Section VIII, B.

**Figure 3:**
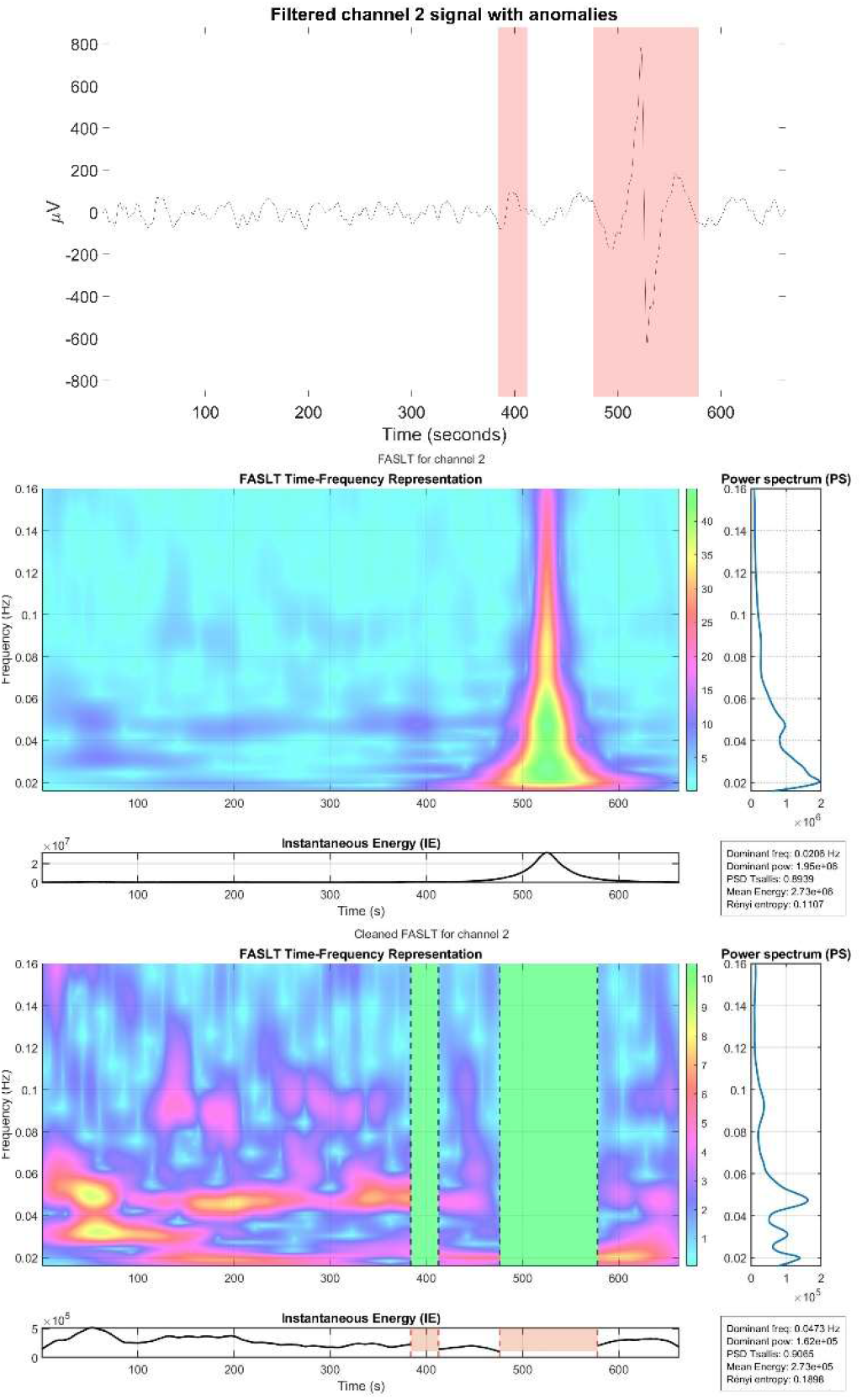
(TOP) Raw EGG signal filtered within the gastric range (0.016– 0.16 Hz). Red bands indicate intervals marked as anomalies. (MIDDLE) Raw TFR corresponding to the signal in the top panel, clearly affected by an artifact at about 530 seconds. (BOTTOM) Results of the cleaning procedure: a normogastric peak emerges in the power spectrum.

To quantitatively evaluate the GSPT impact, in the conservative and non-conservative version, vs a traditional Hampel filter and LMMSE, we computed the validation measures described in section II, D – following the LMMSE correction to identify the false negatives for each approach. Specifically, LMMSE found 198 False Negative events, implying that the frequency content of these events was above 0.16 Hz. These labels were used to move the False Negatives of the other approaches that overlapped with the LMMSE ones to True Negatives. The results of the comparison are represented in Fig 4 where the FN and recall are not displayed for LMMSE, since this method is used to identify the TN. At first glance, the non-conservative version of GSPT was able to identify a larger number of True Positive events than the other methods. On the other hand, the standard Hampel filter identifies a considerably larger number of False Positives, while only a few False Positives were found by the conservative GSPT. Therefore, the conservative GSPT yielded the largest precision, mainly driven by the low FP number, while the non-conservative GSPT version showed the largest recall, driven by the larger TP number.

**Figure 4:**
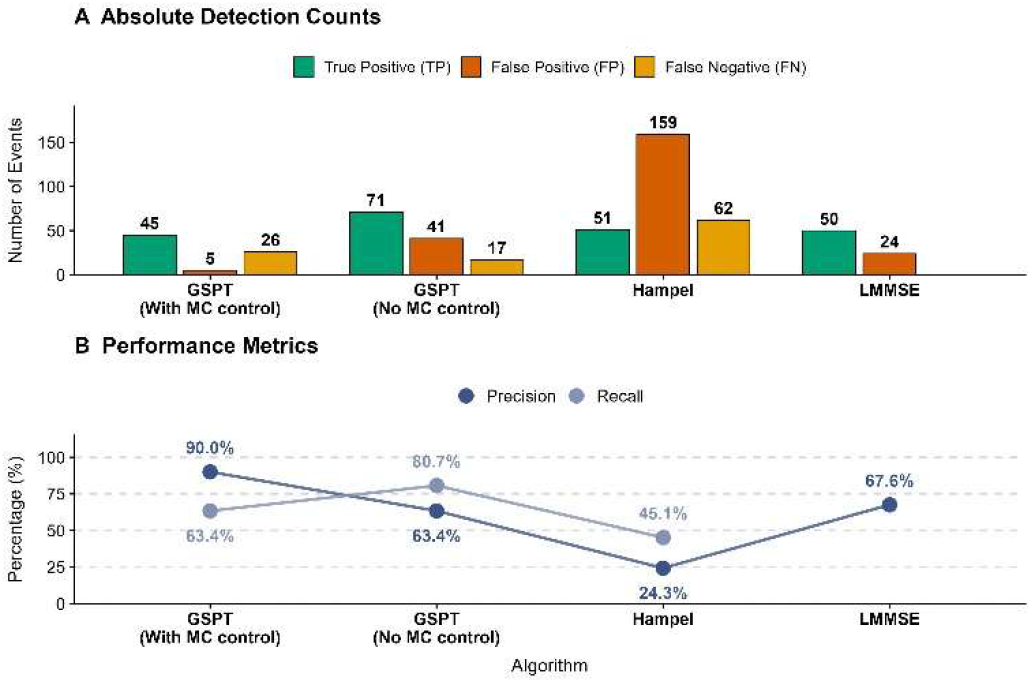
Performance comparison of various algorithms in artifact localization. The stacked bars represent the relative proportion of True Positives (TP, green), False Positives (FP, orange) and False Negatives (FN, yellow) out of the total detected events for each algorithm. For LMMSE, no FN are displayed since this algorithm is used to adjust the FN labeling of the other approaches.

To complement this analysis, we also compared the effect of these different approaches on the dominant peak prominence. GSPT processing—both conservative and non-conservative— yielded a significantly higher peak-to-mean background power ratio compared to both the raw data (with MC: p = 0.02; without MC: p = 0.0002, FDR corrected) and the LMMSE-processed data (with MC: p = 0.02; without MC: p = 0.0002, FDR corrected). Furthermore, the non-conservative GSPT significantly outperformed the Hampel filter (p = 0.0002, FDR corrected). Conversely, no significant differences were observed between Raw and LMMSE, Hampel and Raw, Hampel and LMMSE, and conservative GSPT with Hampel. These results were derived using a cluster bootstrap approach with 10,000 resamples to account for the hierarchical structure of the data (i.e., intra-subject channel correlations). (See Fig S7, Section VIII, C, Supplementary materials).

### D. GSPT OUTCOMES

In the recordings from the 61 subjects, GSPT detected 3.05 ± 2.97 anomalies per recording, with a mean contamination fraction with respect to the total original length of 0.10 ± 0.10; 202 recordings were entirely anomaly-free. Statistical analysis confirmed that noise distribution was spatially uniform: neither the number of anomalies (Fisher’s exact test, p=0.757) nor the fraction of contaminated signal (Kruskal-Wallis, χ^2^(3)=4.42, p=0.220) differed significantly across the four channels.

Two examples of fingerprints from two different channels from two different subjects are shown in Fig. 5. It can be observed that the fingerprints in the first row, extracted from channels 1 and 3 of the same subject, are more similar to each other compared to the fingerprints in the second row, which are extracted from the same two channels of another subject, and vice versa. Qualitatively, fingerprints from the same subject are more coherent with each other compared to those from different subjects.

**Figure 5:**
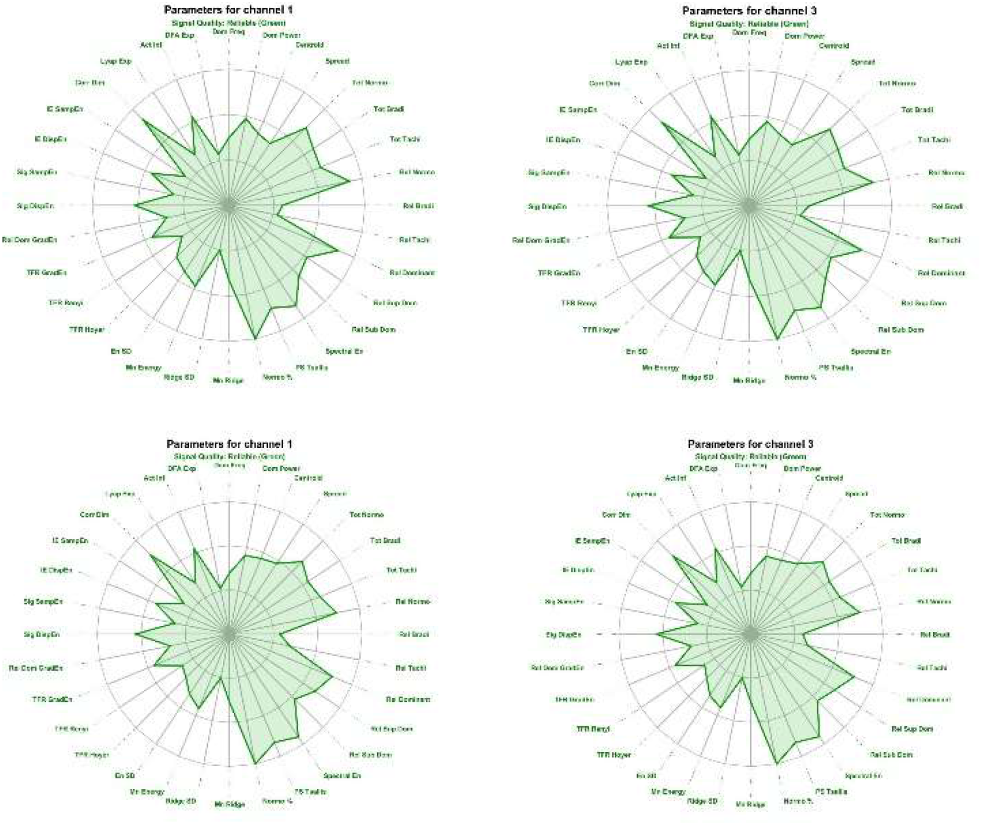
EGG fingerprints from two subjects, obtained from two EGG channels. Clockwise from top left: Subject 1, channel 1; Subject 1, channel 3; Subject 2, channel 3; Subject 2, channel 1.

At a quantitative level, a participant-block bootstrap analysis, with Holm correction across the three planned contrasts, showed the same hierarchy for both fingerprints: Inter-channel similarity> Intra-subject similarity > Inter-subject similarity (see Fig. S8 Section VIII, D, Supplementary materials). For the full fingerprint, the mean correlation advantages were 0.409 (95% bootstrap CI [0.350, 0.467]) for Inter-channel versus Inter-subject, 0.210 [0.144, 0.275] for Intra-subject versus Inter-subject, and 0.199 [0.137, 0.265] for Inter-channel versus Intra-subject (all Holm-adjusted p = 0.0003). The minimal fingerprint reproduced all three contrasts: 0.405 [0.318, 0.490] (p = 0.0003), 0.219 [0.104, 0.333] (p = 0.0006), and 0.186 [0.075, 0.305] (p = 0.0020), respectively. Importantly, the standardised Intra-subject–Inter-subject separation was greater for the full fingerprint (1.201, 95% CI [0.789, 1.720]) than for the minimal fingerprint (0.626 [0.284, 1.052]; difference = 0.575 [0.116, 1.063], p = 0.0161).

These results demonstrate that while factors like electrode placement and daily physiological shifts introduce variability, the individual EGG fingerprint is stable over runs in the same subject and is less similar across subjects. In this light, future studies should assess if EGG fingerprinting might help to distinguish gastric patterns across experimental conditions or between healthy and pathological states.

Furthermore, GSPT estimates instantaneous frequency, phase and amplitude for each recording. An example showing the instantaneous frequency for two different subjects is reported in Fig. 6. Notably, a strong inter-channel similarity within the same individual can be appreciated.

**Figure 6:**
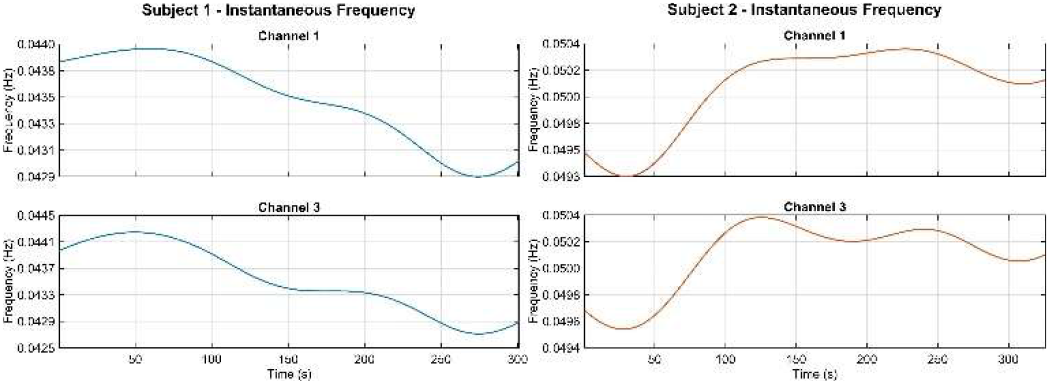
Instantaneous frequency curves for channels 1 and 3, derived from the same subjects in Figure 5.

Finally, we evaluated the cross-session consistency of the nine-feature multivariate fingerprint (Total correlations and multivariate entropies) derived from all four EGG channels. A participant-block bootstrap with 10,000 repetitions showed that Intra-subject correlations (mean r = 0.269, BCa 95% CI [0.156, 0.371]) were higher than Inter-subject correlations (mean r = −0.002, BCa 95% CI [−0.018, 0.032]), with a mean correlation advantage of 0.272 (BCa 95% CI [0.154, 0.378], two-sided bootstrap p < 0.0001).

### E. QUALITY CHECK

The same dataset from 61 subjects was used to evaluate the quality check through MC surrogate tests and the SD consistency selection.

To evaluate the impact of the noise-rejection framework and the quality check, we compared the proportion of DFs falling within the three known physiological bands—bradygastria (<0.033 Hz), normogastria (0.033–0.067 Hz), and tachygastria (>0.067 Hz)—before and after removing channels, following artifact rejection, identified as unreliable under each criterion: MC alone, DF consistency alone, and the combination of MC and DF consistency, with the additional requirement that only recordings retaining at least three channels passed this check. Results are shown in Table 1.

**Table 1:** Effects of the selection methods on the original data. The DF consistency selection considered here uses the Strict threshold (0.005 Hz). Percentages in parentheses are computed relative to the total number of channels retained at each stage (i.e., the row total), not to the original dataset size. The last row is obtained by combining the MC and DF consistency methods, additionally requiring that a recording comprise at least three good channels.

| **Method** | **Bradygastria** | **Normogastria** | **Tachygastria** | **Total** |
|---|---|---|---|---|
| **Raw** | 74 (15.2 %) | 373 (76.4 %) | 41 (8.4 %) | 488 |
| **MC** | 34 (9.3 %) | 313 (85.3 %) | 20 (5.4%) | 367 |
| **DF consistency** | 26 (7.3 %) | 323 (90.5 %) | 8 (2.2 %) | 357 |
| **MC & DF consistency & >=3 ch** | 3 (1.1 %) | 256 (96.2 %) | 7 (2.6 %) | 266 |

First, to evaluate the effect of the MC test, we analyzed the retention rates for each frequency band. A Pearson’s Chi-squared test of independence revealed a highly significant association between the original EGG band and the probability of being retained (χ^2^ (2, N = 488) = 71.26, p < 0.001).

Specifically, the retention rate for normogastric DFs was 84.32%, which was significantly higher compared to both bradygastric DFs (44.15%; post-hoc two-proportion test, FDR corrected: p < 0.001) and tachygastric DFs (48.78%; p < 0.001). The difference in retention rates between bradygastric and tachygastric signals was not statistically significant (p = 0.77). Moreover, we fitted a non-parametric mixed ART model:

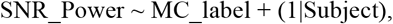

where SNR_Power is the ratio between the mean power in the dominant band (defined as a ±0.015 Hz window around the DF) and the background band (the remaining power within the gastric range). The ‘MC_label’ factor classifies traces based on the Monte Carlo test outcome: ‘Included’ for traces that statistically differed from surrogates, and ‘Excluded’ for the others. We found a significant effect of MC_label (F(1, 472.78) = 218.14, p < 0.001), with the Included traces exhibiting higher values compared to the Excluded ones (Fig S9, Section VIII, E, Supplementary materials). Notably, the spectral skewness and Hoyer measure evaluated by the MC test operate on PSD without any direct link to SNR, by excluding signals lacking the temporally coordinated activity required to generate a distinct peak in the PSD, that is typical of pure noise.

Then, we evaluated the impact of DF consistency for the three proposed thresholds. Specifically, as for the MC test, the percentage of normogastric DFs increased compared to the original condition, while the percentages of bradygastric and tachigastric DF decreased (see Fig. S10, Section VIII, E, Supplementary materials and Table 1 only for the strict threshold). For the strict threshold in this dataset obtained from 4-channel recordings, there are 69 runs that show full retention of all 4 channels, and 28 runs that were recovered only after the elimination of one noisy channel. This suggests that the recursive channel selection procedure is useful to recover recordings that would otherwise be lost.

The specific retention rates per channel for each condition and gastric band are detailed in Fig S11, section VIII E in Supplementary Materials. For each threshold level, we performed a χ^2^ test of independence to assess the association between the original EGG band and the probability of being retained, followed by post-hoc pairwise two-proportion χ^2^ tests with False Discovery Rate (FDR) correction. Across all three threshold conditions, the tests revealed a highly significant association between the gastric band and the retention rate (all p < 0.001). Post-hoc analyses confirmed that the retention of normogastric DFs was significantly higher than that of both bradygastric and tachygastric DFs under all threshold levels (all FDR-corrected p < 0.001), while no significant differences was found between bradygastric and tachygastric signals.

Moreover, mirroring the ART model analysis for the MC-based selection, the signal-to-noise power ratio was significantly higher for Included compared to Excluded signals under the Strict consistency selection criterion (F(1, 476.8) = 53.44, p < 0.001).

Finally, the combined test (last row in Table 1) removed a larger number of channels compared to the single tests, as expected. There are still a few surviving bradygastric and tachygastric DFs. These belong to three subjects, each showing a highly coherent PSD peak shared across three or four channels. The full TFR, along with the PSD and instantaneous energy curve for these three recordings, is reported in Fig. S12, Fig. S13 and Fig. S14 Section VIII, F in the Supplementary Materials. According to GSPT criteria, that allow the analysis of normal and pathological gastric activities, without any a-priori assumption, we cannot exclude that these could be real electrophysiological signals.

## IV. DISCUSSION

While standardized analytical frameworks are well-established for electrophysiological signals such as EEG and ECG, EGG has historically lacked comparable methodological consensus. As a step toward bridging this gap, we developed a novel EGG pipeline designed to facilitate a standardized and automated analysis. EGG presents unique challenges: unlike EEG, it utilizes limited channels, making it difficult to use multichannel information to identify and remove artifacts that are considerably variable over recordings; unlike ECG, it lacks a well-defined morphological pattern, it is an order of magnitude weaker, and is highly susceptible to muscular noise, in addition to electronic noise. To address these challenges, GSPT prioritizes data-driven methodologies applied at the single channel level, and the preservation of signal integrity, that is critical especially during prolonged artifacts. GSPT relies on the Time-Frequency Representation (TFR), that facilitates artifact detection and allows the estimation of several features describing gastric dynamics.

This multidimensional perspective could be essential in clinical and experimental scenarios, such as induction of nausea or disgust, where signals exhibit non-stationary irregularities with multiple spectral peaks and conventional measures such as dominant frequency may, when considered in isolation, provide an incomplete or ambiguous description of gastric activity [5, 34]. Following removal of artifactual intervals, GSPT aims to address this issue by leveraging the full time-frequency topology to introduce complexity-based descriptors, such as sample entropy, which has recently been shown to discriminate nausea-related states from normal gastric activity [35]. Moreover, GSPT provides as output a set of features forming a channel-level fingerprint and a set of instantaneous quantities (frequency, amplitude and phase), that could be potentially used to compare experimental conditions or subjects. Notably, our analyses provided group-level evidence of cross-session participant-specific structure in resting-state EGG fingerprints, as Intra-subject correlations exceeded Inter-subject correlations. Although the same pattern was observed for a minimal fingerprint comprising dominant frequency, dominant power, and fraction of normogastria, the full 32-feature fingerprint produced significantly greater standardised separation between the Intra-subject and Inter-subject distributions. This suggests that the extended fingerprint captures participant-specific multivariate organisation more distinctly than the conventional three-feature representation tested here.

A distinctive feature of GSPT is the preservation of signal integrity in the intervals not affected by artifacts. This philosophy guided the selection of Robust Random Cut Forest and Connectivity Outlier Factor to identify the artefactual epochs, which improve upon Isolation Forest (IF) and Local Outlier Factor (LOF) algorithms in terms of robustness and efficiency. While neural network-based approaches seem promising [30], they remain at the proof-of-concept stage for EGG analysis. Furthermore, beyond the EGG domain, recent large-scale benchmarks in TSAD indicate that deep learning-based methods do not consistently outperform more traditional yet sophisticated algorithms, such as IF or LOF [16].

Crucially, our validation approach, utilizing controlled artifacts and segment-wise metrics offers a quantitative method for assessing algorithm efficacy in artifact localization, aligning with current standards in time-series anomaly detection. This is particularly relevant because the removal or mitigation of movement artifacts could result, in some cases, in small changes to the PSD shape or the DF, which is one of the most widely used parameters in EGG research. Such minor shifts in the DF could be critical, especially when analyzing the effect of different experimental paradigms.

To evaluate the efficacy of GSPT, we quantitatively compared it with the traditional Hampel filter and one of the leading methods in EGG analysis based on the LMMSE estimator.

First, GSPT clearly outperforms the Hampel filter in localizing movement or spike-like artifacts, in terms of precision and recall. Specifically, although the Hampel filter identified a reasonable number of anomalies (similarly to the conservative GSPT and lower than the less conservative GSPT), its performance is severely affected by a high number of false positives and false negatives, achieving low precision and recall compared to GSPT. On the other hand, the conservative GSPT achieved better precision than LMMSE. while the precision of the less conservative GSPT was comparable to LMMSE. However, recall was larger for the less conservative GSPT. Moreover, following artifact removal, both versions of the full GSPT pipeline significantly increased the prominence of the DF relative to the background compared to both raw data and data processed with LMMSE. This suggests that movement artifacts can severely compromise DF quality and that GSPT may be a valuable support in improving DF identification and evaluation. However, it should be noted that while GSPT adopts a signal-rejection (cutting) approach, the LMMSE algorithm is designed to reconstruct and preserve the entire EGG signal. In fact, GSPT is explicitly designed to handle movement or spike-like artifacts while leaving uncontaminated time intervals unaffected, thereby adopting a binary temporal-masking strategy. In contrast, LMMSE follows a more gradual approach: while it primarily targets intervals with major artifacts, it also attenuates less prominent anomalous variations to a lesser degree. Consequently, GSPT and LMMSE represent two distinct methodologies based on different approaches to EGG signal processing.

Furthermore, as a support to standardization of results, a channel-level quality check is applied. The need for standardized, data-driven methodologies led us to introduce two automated procedures for evaluating EGG signal quality, an approach that, to the best of our knowledge, is novel in the EGG field. These two methods, based on IAAFWT MC and DF consistency respectively, have distinct theoretical premises and operate at different scales: the MC is an absolute test performed on each individual channel independently, whereas the DF consistency evaluates all the channels together. These different approaches are generally not independent; specifically as the number of MC-significant channels per recording increased from 0 to 4, the cross-channel standard deviation of dominant frequency tended to decrease [Spearman’s (r_s=-0.511), 95% subject-cluster bootstrap CI [−0.636, −0.358]; two-sided whole-participant permutation (p<0.0001)]. However, while IAAFWT MC maintains its theoretical consistency even in situations where a shift from normogastria is expected, between-channel differences in dominant frequency cannot be attributed exclusively to measurement noise, because impaired spatial coupling and channel-dependent differences in slow-wave frequency and regularity have been reported in functional dyspepsia and systemic sclerosis. [31, 32, 33]. Despite this limitation, DF consistency presumably represents a powerful method that improves upon and automates current state of the art human-based recording selection [4].

Finally, although they were not included in the single-channel fingerprint, GSPT provides two additional classes of measures: (i) single-channel instantaneous phase, frequency, and amplitude and (ii) multichannel total-correlation measures and multivariate entropies (see Fig S15 and Fig S16 in section IX, Supplementary materials for a plot of these GSPT outputs). The latter ones quantify statistical dependences across gastric signals recorded by the entire electrode array. In the present study, the resulting nine-feature multivariate fingerprint showed higher intra-subject correlations across sessions than inter-subject correlations, suggesting that multichannel gastric dependence patterns contain partially reproducible participant-specific information.

## V. METHODOLOGICAL CONSIDERATIONS

While the present results support the effectiveness of GSPT’s artifact-rejection framework, further experimental validation could broaden and strengthen the characterization of EGG artifacts. Concurrent electromyography (EMG) might provide an independent measure of muscular activity; however, establishing its correspondence with EGG artifacts and determining when such activity produces meaningful distortion within the gastric frequency range remains non-trivial. Owing to GSPT’s modular architecture, such extensions could be implemented without disrupting the core workflow, offering opportunities for further refinement and performance optimization.

Future work could integrate GSPT with established multichannel source-separation methods, such as Independent Component Analysis, and explore hybrid strategies combining channel-wise artifact detection with multivariate signal recovery. Such developments could preserve the advantages of channel-specific processing while exploiting shared information across channels, particularly in high-density EGG recordings [36, 37].

A broader limitation for the EGG field is the scarcity of publicly available, validated datasets covering both healthy and clinical populations and containing independently confirmed gastric patterns, such as bradygastria and tachygastria. This constrained our ability to evaluate GSPT in specific physiological and pathological scenarios. Addressing this gap will be important for systematic benchmarking and clinical translation.

## VI. CONCLUSIONS

This work presents an integrated MATLAB framework for automated preprocessing, artifact detection, signal reconstruction, quality assessment, feature extraction, fingerprinting, all at the single-channel level, for the analysis of EGG recordings. Controlled-artifact benchmarks supported the performance of its combined detection strategy for the movement and spike-like perturbations examined, enabling the identification and treatment of contaminated signal segments. Notably, GSPT extracts multiple spectral, time–frequency, instantaneous, and complexity-related descriptors, allowing gastric dynamics to be characterized across complementary dimensions.

GSPT can be exploited to multiple channels, providing in this case an additional fingerprint formed by multi-channel features. Cross-session analyses provided group-level evidence that both single-channel and multichannel fingerprints retain participant-specific structure over time, suggesting possible future exploitation of GSPT to identify experimental conditions or pathologies.

Rather than a static software tool, GSPT is conceived as an extensible methodological framework that can be progressively refined as analytical methods and empirical knowledge in EGG research evolve. Its modular architecture enables new procedures to be incorporated and evaluated without redesigning the core pipeline, while its open-source implementation and native support for an arbitrary number of channels facilitate its application across diverse experimental configurations. The complete workflow can be run through a single execution step, limiting manual intervention and helping to reduce technical barriers to advanced EGG analysis. By combining methodological breadth, adaptability, and accessibility, GSPT aims to make a meaningful contribution to ongoing efforts to improve the robustness, reproducibility, and comparability of EGG research across experimental and clinical settings.

## VII. CODE AVAILABILITY

The GSPT source code is freely available under the MIT License at https://github.com/AleIan2025/Gastric_Signal_Processing_Toolbox---GSPT-

## VIII. CREDIT AUTHORSHIP CONTRIBUTION STATEMENT

**Alessandro Iannone**: Writing – review & editing, Writing – original draft, Visualization, Validation, Software, Methodology, Investigation, Formal analysis, Data curation, Conceptualization. **Maria Serena Panasiti**: Writing – review & editing, Resources. **Salvatore Maria Aglioti**: Writing – review & editing, Project administration, Funding acquisition, Conceptualization. **Stefania Della Penna**: Writing – review & editing, Writing – original draft, Supervision, Methodology, Conceptualization.

## IX. ACKNOWLEDGEMENTS

This work was supported by the European Research Council (ERC) under the European Union’s Horizon 2020 research and innovation programme (eHONESTY, Advanced Grant, grant agreement No. 789058) and Horizon Europe programme (DeepDive, Proof of Concept Grant, grant agreement No. 101138232), and by the Italian Ministry of Health through the Ricerca Finalizzata programme (grant No. GR-2021-12372815).

## BIBLIOGRAPHY

[1] Araki, N., Yamanaka, Y., Poudel, A., Fujinuma, Y., Katagiri, A., Kuwabara, S., & Asahina, M. (2021). Electrogastrography for diagnosis of early-stage Parkinson’s disease. Parkinsonism & Related Disorders, 86, 61–66.

[2] Peralta-Palmezano, J. J., Escobar-Serna, D. P., Peralta-Palmezano, F. J., Acosta-Murillo, N. R., & Guerrero-Lozano, R. (2025). Electrogastrography in pediatric gastroparesis: a systematic review and meta-analysis. Arquivos de Gastroenterologia, 62, e25015.

[3] Al Kafee, A., Cilaci, T., Kayar, Y., & Akan, A. (2022). Electrogastrography in patients with functional dyspepsia, joint hypermobility, and diabetic gastroparesis. The Turkish Journal of Gastroenterology, 33(3), 182.

[4] Wolpert, N., Rebollo, I., & Tallon-Baudry, C. (2020). Electrogastrography for psychophysiological research: Practical considerations, analysis pipeline, and normative data in a large sample. Psychophysiology, 57(9), e13599.

[5] Alladin, S. N., Judson, R., Whittaker, P., Attwood, A. S., & Dalmaijer, E. S. (2024). Review of the gastric physiology of disgust: Proto-nausea as an under-explored facet of the gut– brain axis. Brain and Neuroscience Advances, 8, 23982128241305890.

[6] Richter, C. G., Babo-Rebelo, M., Schwartz, D., & Tallon-Baudry, C. (2017). Phase-amplitude coupling at the organism level: The amplitude of spontaneous alpha rhythm fluctuations varies with the phase of the infra-slow gastric basal rhythm. NeuroImage, 146, 951–958.

[7] Rebollo, I., Devauchelle, A. D., Béranger, B., & Tallon-Baudry, C. (2018). Stomach-brain synchrony reveals a novel, delayed-connectivity resting-state network in humans. elife, 7, e33321.

[8] Komorowski, D., & Mika, B. (2021). Gastric slow wave rhythm identification using new approach based on noise-assisted multivariate empirical mode decomposition and Hilbert–Huang transform. Neurogastroenterology & Motility, 33(3), e13997.

[9] Stallone, A., Cicone, A., & Materassi, M. (2020). New insights and best practices for the successful use of Empirical Mode Decomposition, Iterative Filtering and derived algorithms. Scientific reports, 10(1), 15161.

[10] Gharibans, A. A., Smarr, B. L., Kunkel, D. C., Kriegsfeld, L. J., Mousa, H. M., & Coleman, T. P. (2018). Artifact rejection methodology enables continuous, noninvasive measurement of gastric myoelectric activity in ambulatory subjects. Scientific reports, 8(1), 5019.

[11] Moca, V. V., Bârzan, H., Nagy-Dăbâcan, A., & Mureşan, R. C. (2021). Time-frequency super-resolution with superlets. Nature communications, 12(1), 337.

[12] Golyandina, N. (2025). Singular spectrum analysis. In International Encyclopedia of Statistical Science (pp. 2345–2348). Berlin, Heidelberg: Springer Berlin Heidelberg.

[13] Lancaster, G., Iatsenko, D., Pidde, A., Ticcinelli, V., & Stefanovska, A. (2018). Surrogate data for hypothesis testing of physical systems. Physics reports, 748, 1–60.

[14] Tang, J., Chen, Z., Fu, A. W. C., & Cheung, D. W. (2002, April). Enhancing effectiveness of outlier detections for low density patterns. In Pacific-Asia conference on knowledge discovery and data mining (pp. 535–548). Berlin, Heidelberg: Springer Berlin Heidelberg.

[15] Guha, S., Mishra, N., Roy, G., & Schrijvers, O. (2016, June). Robust random cut forest based anomaly detection on streams. In International conference on machine learning (pp. 2712–2721). PMLR.

[16] Boniol, P., Paparrizos, J., Kang, Y., Palpanas, T., Tsay, R. S., Elmore, A. J., & Franklin, M. J. (2022). Theseus: Navigating the Labyrinth of Time-Series Anomaly Detection. Proc. VLDB Endow., 15(12), 3702–3705.

[17] Hundman, K., Constantinou, V., Laporte, C., Colwell, I., & Soderstrom, T. (2018, July). Detecting spacecraft anomalies using lstms and nonparametric dynamic thresholding. In Proceedings of the 24th ACM SIGKDD international conference on knowledge discovery & data mining (pp. 387–395).

[18] Brevdo, E. & Thakur, G. MATLAB Synchrosqueezing Toolbox. GitHub https://github.com/ebrevdo/synchrosqueezing (2011).

[19] Thakur, G., Brevdo, E., Fučkar, N. S., & Wu, H. T. (2013). The Synchrosqueezing algorithm for time-varying spectral analysis: robustness properties and new applications. Signal Processing, 93(5), 1079–1094.

[20] Weber, I., & Oehrn, C. R. (2022). NOLITIA: an open-source toolbox for non-linear time series analysis. Frontiers in neuroinformatics, 16, 876012.

[21] Iatsenko, D., McClintock, P. V., & Stefanovska, A. (2016). Extraction of instantaneous frequencies from ridges in time–frequency representations of signals. SignalProcessing, 125, 290–303.

[22] Flood, M. W., & Grimm, B. (2021). EntropyHub: An open-source toolkit for entropic time series analysis. PloS one, 16(11), e0259448.

[23] Watanabe, S. (1960). Information theoretical analysis of multivariate correlation. IBM Journal of research and development, 4(1), 66–82.

[24] Giraldo, L. G. S., Rao, M., & Principe, J. C. (2014). Measures of entropy from data using infinitely divisible kernels. IEEE Transactions on Information Theory, 61(1), 535–548.

[25] Yu, S., Giraldo, L. G. S., Jenssen, R., & Principe, J. C. (2019). Multivariate Extension of Matrix-Based Rényi’s $\alpha $ α-Order Entropy Functional. IEEE transactions on pattern analysis and machine intelligence, 42(11), 2960–2966.

[26] Gharibans, A. A., Coleman, T. P., Mousa, H., & Kunkel, D. C. (2019). Spatial patterns from high-resolution electrogastrography correlate with severity of symptoms in patients with functional dyspepsia and gastroparesis. Clinical Gastroenterology and Hepatology, 17(13), 2668–2677.

[27] Buschow, S. (2023). dualtrees: Decimated and Undecimated 2D Complex Dual-Tree Wavelet Transform (R package version 0.1.5). CRAN. 10.32614/CRAN.package.dualtrees

[28] Simes, R. J. (1986). An improved Bonferroni procedure for multiple tests of significance. Biometrika, 73(3), 751–754.

[29] Montgomery, D. C. (2020). Introduction to statistical quality control. John wiley & sons.

[30] Liang, J., Cheung, J. Y., & Chen, J. D. Z. (1997). Detection and deletion of motion artifacts in electrogastrogram using feature analysis and neural networks. Annals of Biomedical Engineering, 25(5), 850–857

[31] Lin, X., & Chen, J. Z. (2001). Abnormal gastric slow waves in patients with functional dyspepsia assessed by multichannel electrogastrography. American journal of physiology-gastrointestinal and liver physiology, 280(6),G1370–G1375

[32] McNearney, T., Lin, X., Shrestha, J., Lisse, J., & Chen, J.D. (2002). Characterization of gastric myoelectrical rhythms in patients with systemic sclerosis using multichannel surface electrogastrography. Digestive diseases and sciences, 47(4), 690–698.

[33] Sha, W., Pasricha, P. J., & Chen, J. D. (2009). Rhythmic and spatial abnormalities of gastric slow waves in patients with functional dyspepsia. Journal of clinical gastroenterology, 43(2), 123–129.

[34] Carson, D. A., O’Grady, G., Du, P., Gharibans, A. A., & Andrews, C. N. (2021). Body surface mapping of the stomach: New directions for clinically evaluating gastric electrical activity. Neurogastroenterology & Motility, 33(3), e14048

[35] Jakus, G., Sodnik, J., & Miljković, N. (2022). Electrogastrogram-derived features for automated sickness detection in driving simulator. Sensors, 22(22), 8616.

[36] Sengottuvel, S., Khan, P. F., Mariyappa, N., Patel, R., Saipriya, S., & Gireesan, K. J. S. T. T. L. S. I. (2018). A combined methodology to eliminate artifacts in multichannel electrogastrogram based on independent component analysis and ensemble empirical mode decomposition. SLAS TECHNOLOGY: Translating Life Sciences Innovation, 23(3), 269–280.

[37] O’Grady, G., Angeli, T. R., Paskaranandavadivel, N., Erickson, J. C., Wells, C. I., Gharibans, A. A., … & Du, P. (2018). Methods for high-resolution electrical mapping in the gastrointestinal tract. IEEE reviews in biomedical engineering, 12, 287–302.

