## Supplementary Material for "Optimizing non-invasive assessment of gastric activity: a fully automated channel-wise pipeline for multi-electrode electrogastrography"

### SUPPLEMENTARY MATERIALS

#### I. SPECTRAL CONTENT OF JUMPS

Spectral content of the jumps can be estimated modelling the jump as the Heaviside step function H(t). Let's call our recorded signal s(t) and let's assume that there is a jump at a time $t=t^*$, then we can write:

$$s(t) = c(t) + A \cdot H(t - t^*),$$

where c(t) is the clean signal without the jump and A is a constant modelling the amplitude of the jump. If we further suppose that the power spectrum density (PSD) of c(t) is negligible above the Nyquist frequency $f_s/2$, where $f_s$ is the sampling frequency, than the PSD of s(t) above $f_s/2$ is mainly due to AH(t-t*). First of all, let's calculate the Fourier transform of s(t):

$$s\widetilde{(w)} = c\widetilde{(w)} + Ae^{-iwt^*}H\widetilde{(w)}.$$

The PSD of s(t) is the modulus squared of $s\widetilde{(w)}$ and it is composed by three terms:

$$\left|((s))\widetilde{(w)}\right|^2 = \left|((c))\widetilde{(w)}\right|^2 + A^2\left|((H))\widetilde{(w)}\right|^2 + \text{interference}.$$

The first term by hypothesis is negligible above $f_s/2$ and the interference terms are linear in $\widetilde{c(w)}$, so that even these terms is negligible above $f_s/2$. The only relevant term is thus the one containing the Fourier transform of the Heaviside function:

$$((H))\widetilde{(w)} = \pi\delta(w) - \frac{i}{w}.$$

The PSD of the modelled jump at $f_s/2$ is

$$PSD\left(\frac{f_s}{2}\right) = \frac{A^2}{w^2} = \frac{A^2}{\pi^2 f_s^2}.$$

Thus, the jump contribution to the spectrum above the Nyquist frequency is negligible, when

$$A \ll \pi f_s.$$

Notably, this upper limit depends on how we model the jump. IN GSPT, we selected a conservative threshold U= $\frac{\pi f_s}{8}$. We thus compared the absolute value of the discrete derivative of the signal at each time point with this threshold. Any derivative showing a value larger than this threshold should be corrected to avoid aliasing.

Note that with this approach it is also possible to model a spike by combining two Heaviside functions whose steps differ by one sample, or $\delta t = 1/f_s$:

$$s(t) = c(t) + AH(t - t^*) - AH(t - t^* - \delta t).$$

Thus, the power spectrum density at $f_s/2$ is:

$$PSD\left(\frac{f_s}{2}\right) = \frac{2A^2}{w^2}(1 - \cos(w\delta t))|_{w=\pi f_s} = \frac{2A^2}{\pi^2 f_s^2}(1 - \cos(\pi f_s \delta t)) = \frac{4A^2}{\pi^2 f_s^2}.$$

Hence, a threshold like the one for the jumps can be used.

#### II. TSALLIS ENTROPY

Tsallis entropy is a generalization of the classical Shannon entropy, and it is defined by the following formula [1]:

$$TE_q[p] = \frac{1}{q-1}\left(1 - \int (p(x))^q dx\right),$$

where q is a free parameter and p(x) is the probability distribution to which we want to associate TE. In the limit of q tending to 1 this entropy reduces to the Shannon entropy [1].

The key advantage of the Tsallis entropy over the classical Shannon entropy lies in the tunable parameter q. Compared to Shannon entropy (q=1), Tsallis entropy with q>1 downweights rare events (e.g., spikes corresponding to high p(x) values), while q<1 amplifies their contribution. Since our goal is spike detection, Tsallis entropy with q<1 provides superior sensitivity to transient artifacts compared to Shannon entropy.

In our framework, we estimate TE of normalized Instantaneous Energy (IE) curve, scaled to have an integral equal to one. A flat IE curve corresponds to high entropy, whereas a sharply peaked IE curve (indicating signal concentration) exhibits low entropy. To optimize discriminative power, we compute Tsallis entropy at the q value corresponding to the minimum entropy for each signal. This approach not only maximizes artifact detectability but also adaptively selects the optimal q for each unique IE curve.

$$TE_{q_{min}}[IE] = \frac{1}{q_{min} - 1}\left(1 - \int \left(\widetilde{IE}(x)\right)^{q_{min}} dx\right)$$

Figure S1 displays an example of a signal affected by strong artifacts. The related TE (0.6752) is significantly smaller than the MC surrogate distribution (0.8891±0.0169).

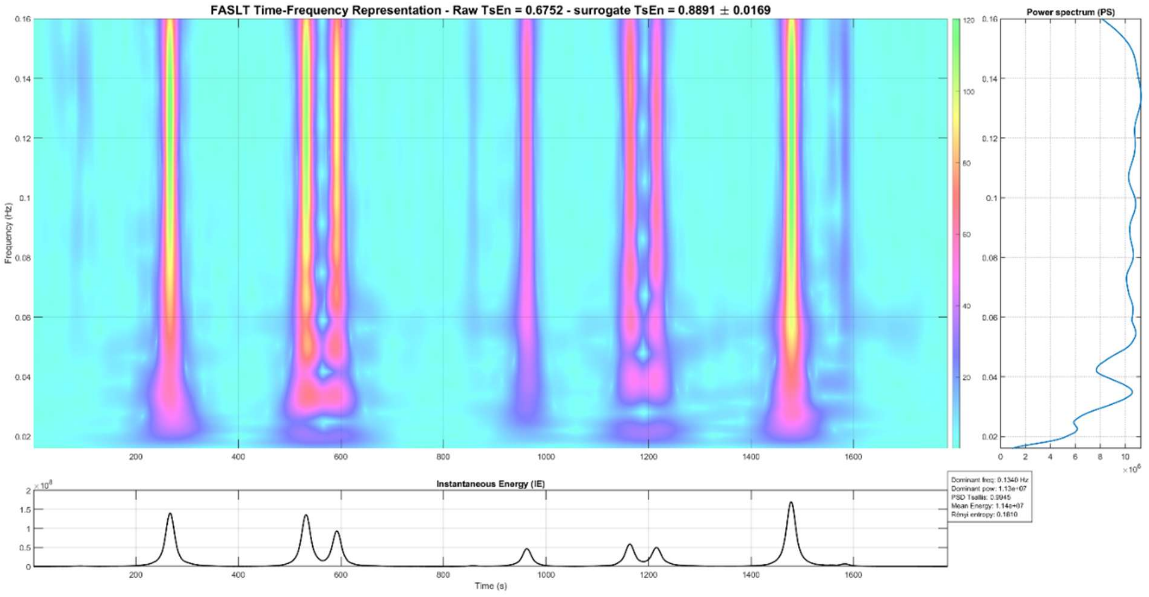


Figure S1: A TFR for a real recorded EGG signal with strong artifacts. The IE clearly shows extreme spikes.: The signal's IE TE is significantly smaller than the result of the surrogate test, reflecting the presence of high peaks in the IE curve. Thus, the MC test correctly classifies this signal being contaminated by spike-like artifacts.

### III. TSAD EXAMPLES

A local Tukey fences method analyses the value distribution of the filtered EGG signal using sliding windows of fixed length (100 seconds). Consecutive windows overlap almost entirely, differing by only one data point. For each window, the central point is evaluated against the distribution of values in that window. If this point falls outside the classical 1.5 interquartile range (IQR) threshold, it is flagged as an anomaly; otherwise, it is classified as normal. This iterative process ensures a dynamic and localized assessment of signal integrity. An example of this procedure can be found in Fig S2.

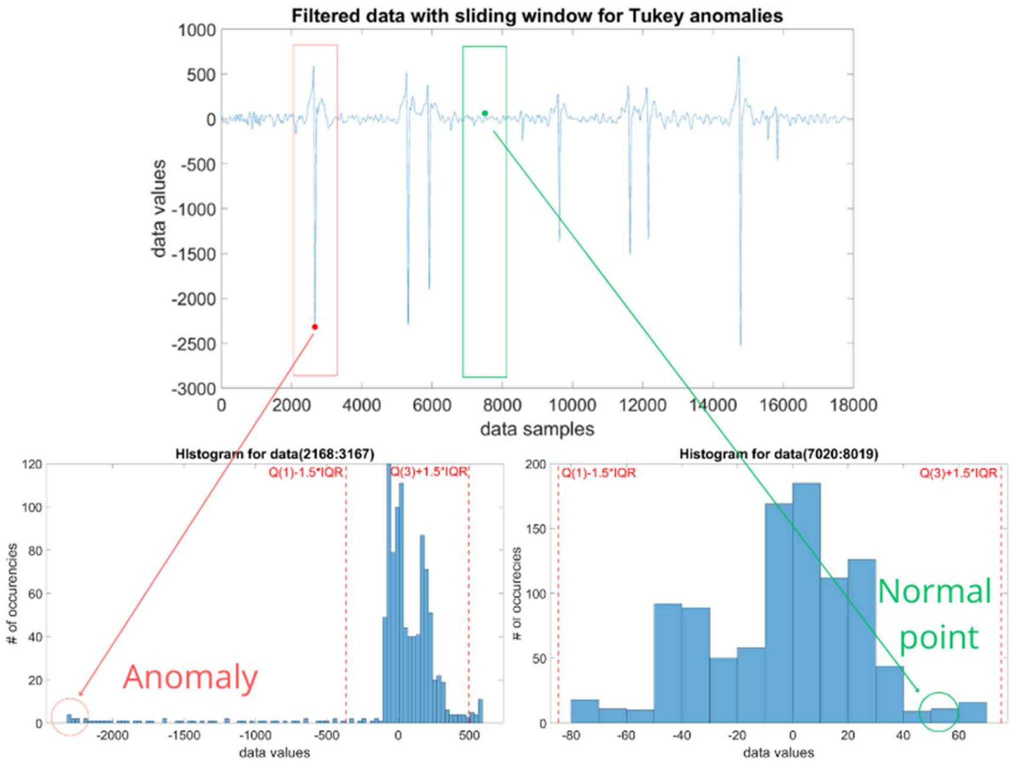


*Figure S2: The top of the figure displays a real EGG signal, filtered in the gastric frequency range. Analyzing the central point of two distinct windows yields contrasting results. The data point within the red window lies far outside the local distribution and is consequently flagged as an anomaly. In contrast, the point in the green window is not extreme, falling below the $Q_3 + 1.5 * IQR$ threshold. Thus, it is classified as a normal point by the Local Tukey Fences algorithm.*

### IV. RELATIVE POWERS DEFINITIONS

In this section, we report the definitions of relative power in the classical frequency bands (normogastria, bradygastria, and tachygastria), along with our adaptive band definition centred around the detected dominant frequency for each signal. For each band, the integral of the power spectrum density is divided by the bandwidth to compute a power per unit frequency, allowing for comparison between bands of different widths.

| Name | Definition | Explanation |
|---|---|---|
| Mean relative normogastric power | $\dfrac{\int_{0.033}^{0.067} \frac{PSD(f)}{0.067-0.033} df}{\int_{0.033}^{0.067} \frac{PSD(f)}{0.067-0.033} df + \int_{0.067}^{0.16} \frac{PSD(f)}{0.16-0.067} df + \int_{0.016}^{0.033} \frac{PSD(f)}{0.033-0.016} df}$ | The ratio of the mean power density in the normogastric band to the sum of the mean power density in each classical frequency band. Its values range from 0 to 1 |

| | | |
|---|---|---|
| Mean relative tachygastric power | $$\frac{\int_{0.067}^{0.16}\frac{PSD(f)}{0.16-0.067}df}{\int_{0.033}^{0.067}\frac{PSD(f)}{0.067-0.033}df+\int_{0.067}^{0.16}\frac{PSD(f)}{0.16-0.067}df+\int_{0.016}^{0.033}\frac{PSD(f)}{0.033-0.016}df}$$ | The ratio of the mean power density in the tachygastric band to the sum of the mean power density in each classical frequency band. Its values range from 0 to 1 |
| Mean relative bradygastric power | $$\frac{\int_{0.016}^{0.033}\frac{PSD(f)}{0.033-0.016}df}{\int_{0.033}^{0.067}\frac{PSD(f)}{0.067-0.033}df+\int_{0.067}^{0.16}\frac{PSD(f)}{0.16-0.067}df+\int_{0.016}^{0.033}\frac{PSD(f)}{0.033-0.016}df}$$ | The ratio of the mean power density in the bradygastric band to the sum of the mean power density in each classical frequency band. Its values range from 0 to 1 |
| Dominant band relative power | $$\frac{\int_{DF-0.015}^{DF+0.015}\frac{PSD(f)}{0.030}df}{\int_{DF+0.015}^{0.16}\frac{PSD(f)}{0.16-(DF+0.015)}df+\int_{0.016}^{DF-0.015}\frac{PSD(f)}{(DF-0.015)-0.016}df}$$ | The relative mean power density of the signal around its dominant frequency respect to the frequency band above and below it |
| Sub-dominant band relative power | $$\frac{\int_{0.016}^{DF-0.015}\frac{PSD(f)}{(DF-0.015)-0.016}df}{\int_{DF-0.015}^{DF+0.015}\frac{PSD(f)}{0.030}df+\int_{DF+0.015}^{0.16}\frac{PSD(f)}{0.16-(DF+0.015)}df}$$ | The relative mean power density of the signal in the frequency band below the dominant frequency band respect to the other two bands |
| Super-dominant band relative power | $$\frac{\int_{DF+0.015}^{0.16}\frac{PSD(f)}{0.16-(DF+0.015)}df}{\int_{DF-0.015}^{DF+0.015}\frac{PSD(f)}{0.030}df+\int_{0.016}^{DF-0.015}\frac{PSD(f)}{(DF-0.015)-0.016}df}$$ | The relative mean power density of the signal in the frequency band above the dominant frequency band respect to the other two bands |

*Table S1: Spectral power ratios definitions. In the formulas, PSD denotes the Power Spectrum Density, DF the Dominant Frequency, and f the frequency. The total gastric range spans from 0.016 to 0.16 Hz. Standard frequency sub-bands are defined as follows:*

*bradygastria (0.016–0.033 Hz), normogastria (0.033–0.067 Hz), and tachygastria (0.067–0.16 Hz). The adaptive bands are instead defined relative to the dominant frequency: the dominant band corresponds to an interval of ±0.015 Hz (capped at the boundaries of the gastric spectrum if the interval exceeds them) centered on the DF. Consequently, the sub- and super-dominant bands comprise the remaining lower and upper frequency ranges within the total gastric spectrum (0.016–0.16 Hz).*

### V. NOVEL QUANTITIES FOR EGG

Table S2 lists metrics that are novel in EGG, defined via mathematical formulas or references to the code implementations, alongside a brief explanation of their meaning.

All quantities requiring a phase space embedding (e.g., dispersion entropy, correlation dimension) depend on the proper estimation of the delay τ and the embedding dimension m. These parameters are estimated using the phaseSpaceReconstruction MATLAB function, which implements the standard False Nearest Neighbors (FNN) and Average Mutual Information (AMI) algorithms.

| **Name** | **Definition/Reference** | **Explanation** |
|---|---|---|
| Spectral centroid | Spectral descriptors, Audio Toolbox, MATLAB [2] | The "centre of mass" of the power spectrum |
| Spectral spread | Spectral descriptors, Audio Toolbox, MATLAB [2] | The spread of the power spectrum around its spectral centroid |
| Spectral entropy | Spectral descriptors, Audio Toolbox, MATLAB [2] | Classical Shannon entropy of the signal power spectrum. The higher the spectral entropy the more diffuse is the power spectrum |
| Tsallis entropy of the Power Spectrum Density (PSD) | $TS_{PSD} = \frac{1}{q_{min}-1}\left(1 - \int_{0.016}^{0.16} \left(PSD_n(f)\right)^{q_{min}} df\right),$ $PSD_n(f) \triangleq PSD(f) / \int_{f_{gastric}} PSD(f)df$ | An entropy with generally higher sensitivity to changes in the PSD respect to classical spectral entropy |
| Mean ridge frequency | The mean of the instantaneous frequency curve extracted directly from the superlet TFR ridges | It reflects the time-averaged trajectory of the signal's dominant frequency, giving equal weight to every temporal instant regardless of its instantaneous power |
| Ridge frequency standard deviation | The standard deviation of the instantaneous frequency curve extracted directly from the superlet TFR | A measure of the overall dispersion of the instantaneous frequency ridge trajectory |
| Mean energy | The mean of the Instantaneous energy curve | The mean energy of the signal computed from its TFR |
| Energy standard deviation | The standard deviation of the instantaneous energy curve | A measure of the overall variability of the IE's values |
| Dispersion entropy estimated from IE | EntropyHub function, [3] | Quantifies the predictability of a signal by measuring how uniformly its values are distributed across symbolic categories. Quantifies the temporal complexity and unpredictability of the IE. It captures dynamic patterns rather than simple magnitude dispersion. |
| Sample entropy estimated from IE | EntropyHub function, [3] | Estimates signal complexity by counting the rarity of repeated patterns, with |

| | | |
|---|---|---|
| | | fewer repetitions indicating higher entropy. Quantifies the temporal complexity and unpredictability of the IE. It captures dynamic patterns rather than simple magnitude dispersion. |
| Dispersion entropy estimated from the raw signal | EntropyHub function, [3] | Same entropy described in the row for the Dispersion entropy estimated from IE, this time computed on the raw signal filtered in the gastric range (0.016-0.16 Hz) |
| Sample entropy estimated from the raw signal | EntropyHub function, [3] | Same entropy described in the row for the Sample entropy estimated from IE, this time computed on the raw signal filtered in the gastric range (0.016-0.16 Hz) |
| TFR Renyi entropy | $R_\alpha = \frac{1}{1-\alpha} \log_2 \left( \iint p(t,f)^\alpha \, df \, dt \right)$<br>$p(t,f) \equiv \frac{\|TFR(t,f)\|^2}{\iint \|TFR(t,f)\|^2 \, df \, dt}$<br>Where TFR(t, f) is the time frequency matrix at time t and frequency f.<br>[4] | A measure of concentration of the TFR. The more the energy of the signal is confined in a small time and frequency space, the more the Renyi entropy is low |
| Hoyer measure | $H_M = \frac{\sqrt{N} - \frac{\\|\mathrm{TFR}\\|_1}{\\|\mathrm{TFR}\\|_2}}{\sqrt{N} - 1}$<br>$\\|\mathrm{TFR}\\|_1 = \iint_\Omega \|\mathrm{TFR}(t,f)\| \, dt \, df$<br>$\\|\mathrm{TFR}\\|_2 = \sqrt{\iint_\Omega \|\mathrm{TFR}(t,f)\|^2 \, dt \, df}$<br>$N = \iint_\Omega dt \, df$<br>[5] | A measure of concentration of the TFR. The more the energy of the signal is confined in a small time and frequency space, the more the Hoyer measure is low |
| Gradient entropy (GradEn) | Matlab code implemented following [6] | Quantifies the irregularity of image textures; here it is applied to the TFR treated as an image |
| Relative gradient entropy | The GradEn within the dominant band divided by the sum of the GradEn values in the adjacent higher and lower frequency bands. | Quantifies the irregularity of image textures. Here, it is applied to the TFR as an image and compares the irregularity in the dominant band with that in the rest of the TFR. |
| Correlation dimension | NoLiTiA function, [7] | A mathematical tool that measures the "geometric complexity" of a dataset by analysing how its points fill a multidimensional space. It distinguishes between |

| | | |
|---|---|---|
| | | systems governed by random noise (simple, low-dimensional patterns) and deterministic chaos (complex, fractal-like behavior). For instance, regular periodic signals (e.g., stable heartbeats) exhibit low dimensionality, while chaotic systems (e.g., turbulent fluid flow) display higher dimensionality. |
| Lyapunov exponent | NoLiTiA function, [7] | Measures the exponential divergence rate of nearby trajectories in a dynamical system. If the exponent is positive, starting from two very close values in the time series led to wildly different behaviour after a small amount of time. If the exponent is negative, close points led to close future trajectories even after a significant time interval |
| Active information storage (AIS) | NoLiTiA function, [7] | Quantifies the amount of information from a system's past that is actively used to influence its present state. High AIS implies strong dependency on historical data (e.g., neurons retaining past activity for decision-making). |
| Detrend fluctuation analysis (DFA) exponent | NoLiTiA function, [7] | A more robust estimate of the hurst exponent. It quantifies long-range correlations in time series and reflects the likelihood of a value increase/decrease being followed by another similar trend.<br>$\alpha>0.5$: Persistence (trends tend to continue, e.g., increases follow increases).<br>$\alpha = 0.5$: Uncorrelated noise (no predictable pattern, akin to random walks).<br>$\alpha < 0.5$: Anti-persistence (trends reverse direction more frequently than random noise). |

*Table S2: Novel single-channel metrics proposed for EGG analysis.*

### VI. EXAMPLE OF NOISE EFFECTS

The power spectrum in Fig S3a suggests a genuine DF in the normogastria range, yet it was derived from a signal recorded by an EGG device with no electrodes attached—thus representing purely electronic noise. This clearly demonstrates how noise alone could mimic physiologically plausible spectral features, highlighting the need for quantitative signal-quality assessment before interpreting EGG-derived parameters. Inspection of the corresponding full

TFR (Fig. S3b) may nevertheless provide useful qualitative insights by revealing temporal and spectral irregularities that are not apparent from the power spectrum alone.

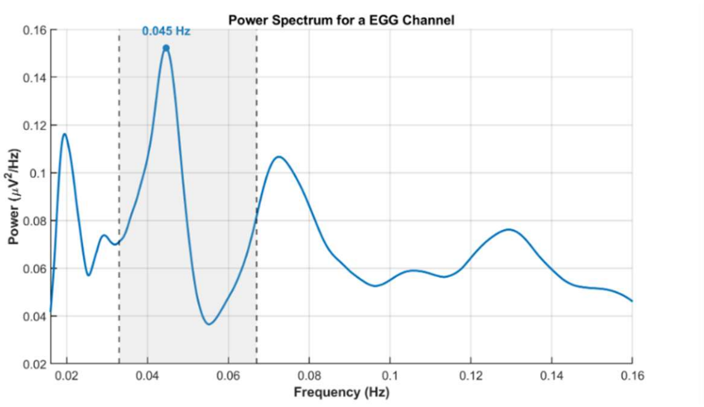


Figure S3a: Power spectrum density for a real EGG recording without electrodes attached

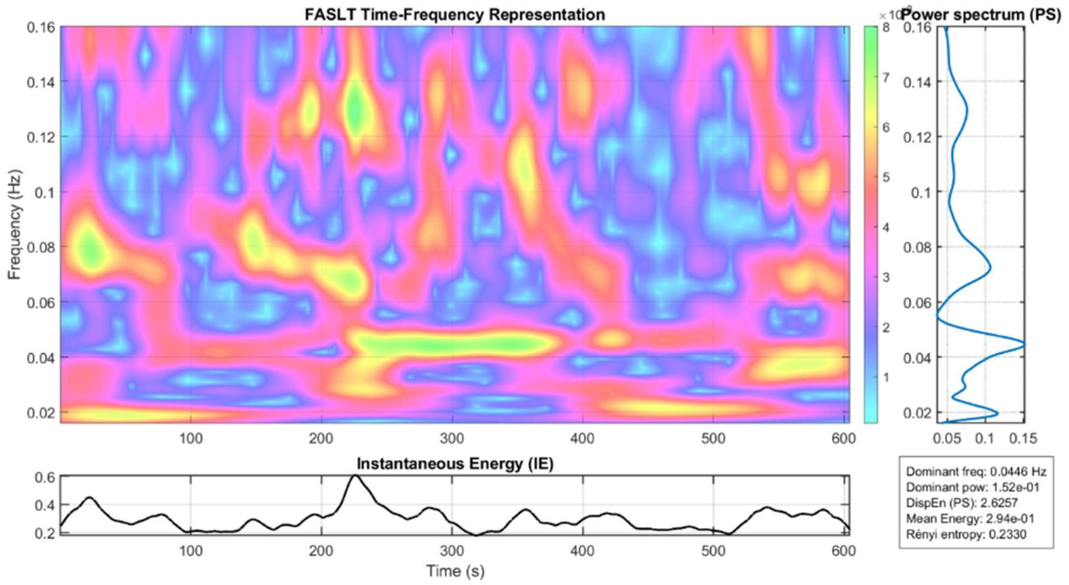


Figure S3b: The full TFR associate to the power spectrum in figure S3a

### VII. SPECTRAL SKEWNESS AND HOYER ENTROPY

If P(x) indicates the power spectrum density and $NP(x) = \frac{P(x)}{\int P(x)dx}$ the normalized power spectrum, the spectral skewness can be defined as follows:

$$\gamma_1 = \frac{\int (x-\mu)^3 \cdot NP(x)\, dx}{\sigma^3}$$

Where $\mu$ and $\sigma$ are the mean and dispersion of $NP(x)$.

Let NF the number of frequency bins of interest (corresponding in our case to [0.016, 0.16] Hz), then the Hoyer entropy [8] is defined by the following formula:

$$Hy(P) = \frac{\sqrt{N_F} - \frac{||P||_1}{||P||_2}}{\sqrt{N_F} - 1}$$

Where

$$||P||_1 = \int P(x)\,dx, \qquad ||P||_2 = \sqrt{\int P(x)^2\,dx}$$

### VIII. RESULTS: SUPPORTING IMAGES AND EXAMPLES

#### A. PREPROCESSING

Here, we report the raincloud representation of the Tsallis entropy on instantaneous energy (Figure S4; see Main Text, Results, Section B).

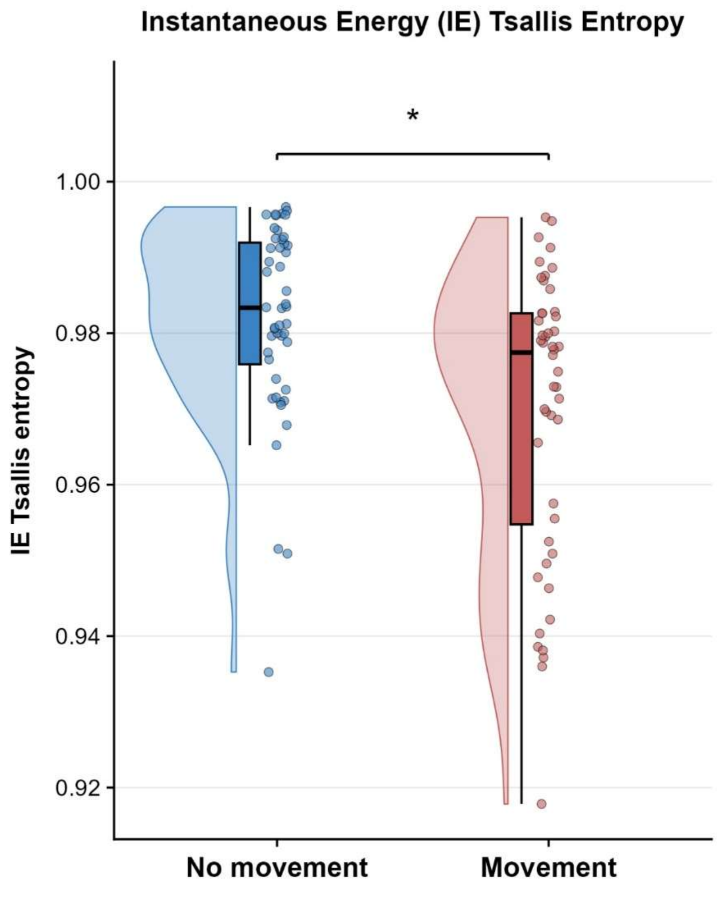


*Figure S4: Tsallis entropy of the instantaneous energy for participants performing controlled movements versus participants at rest.*

### B. EXAMPLES OF TSAD AND TFR CLEANING

*We here show additional examples on GSPT performances.* In Fig. S5, the results of the full TSAD algorithm are shown for two real EGG recordings obtained from two participants with controlled movement artifacts, where anomalous intervals can be easily spotted.

Figures S6a and S6b illustrate the results of the full artifact rejection procedure, combining TSAD and TFR cleaning, applied to two additional real EGG recordings. In the raw data, the dominant frequency is driven by induced artifacts, shifting downwards to approximately 0.03 Hz (Fig. S6a, top panel) or upwards to approximately 0.07 Hz (Fig. S6b, top panel). Following TFR cleaning, the dominant frequency is restored to about 0.05 Hz in Figure S6a (bottom panel) and 0.04 Hz in Figure S6b (bottom panel), while the PSD at the artifactual dominant frequencies is substantially reduced. These examples demonstrate that artifacts can induce spurious bradygastric as well as tachygastric dominant frequencies, and that the proposed cleaning procedure *is capable of handling both scenarios*.

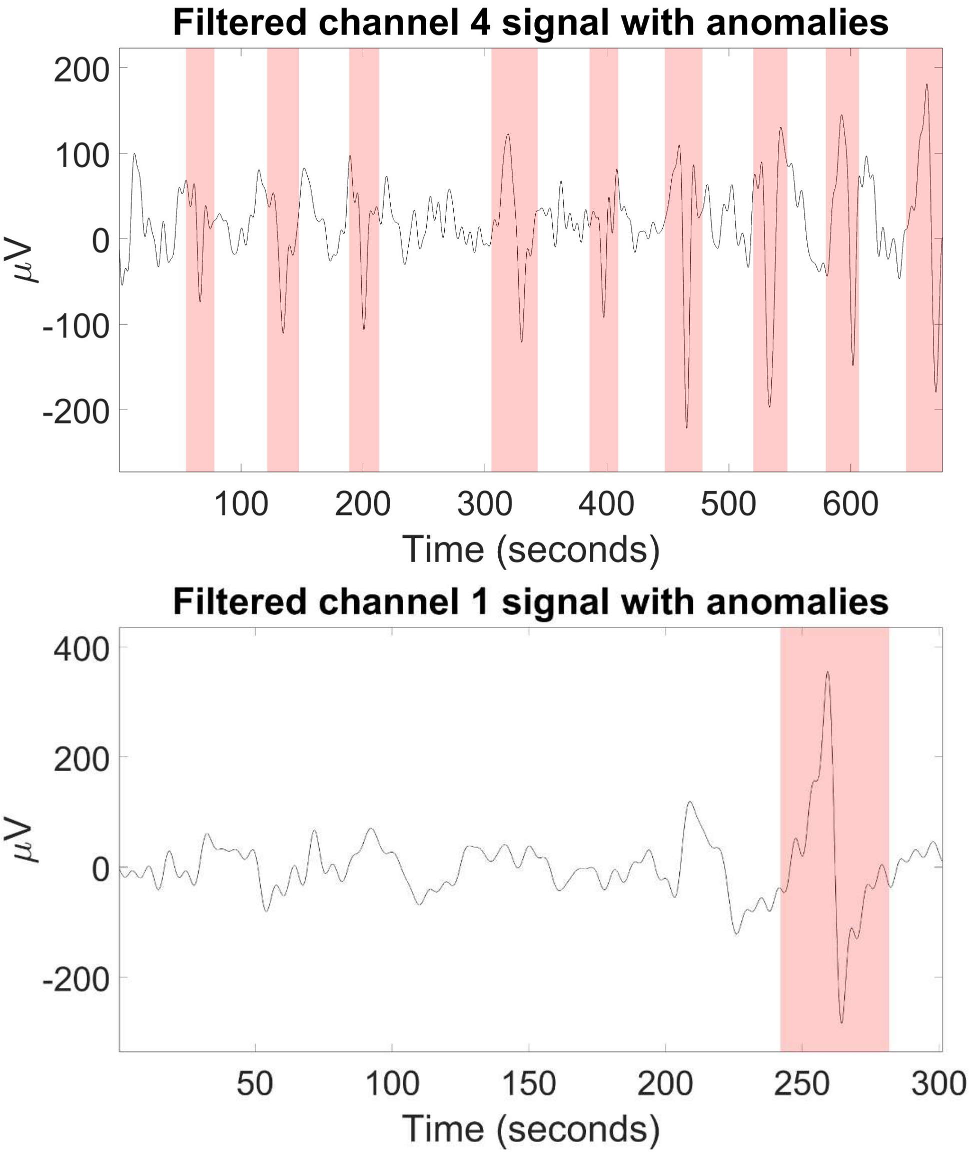


*Figure S5: two recorded signals with controlled anomalies for two different subjects. The ones detected by the algorithm are highlighted with red bands.*

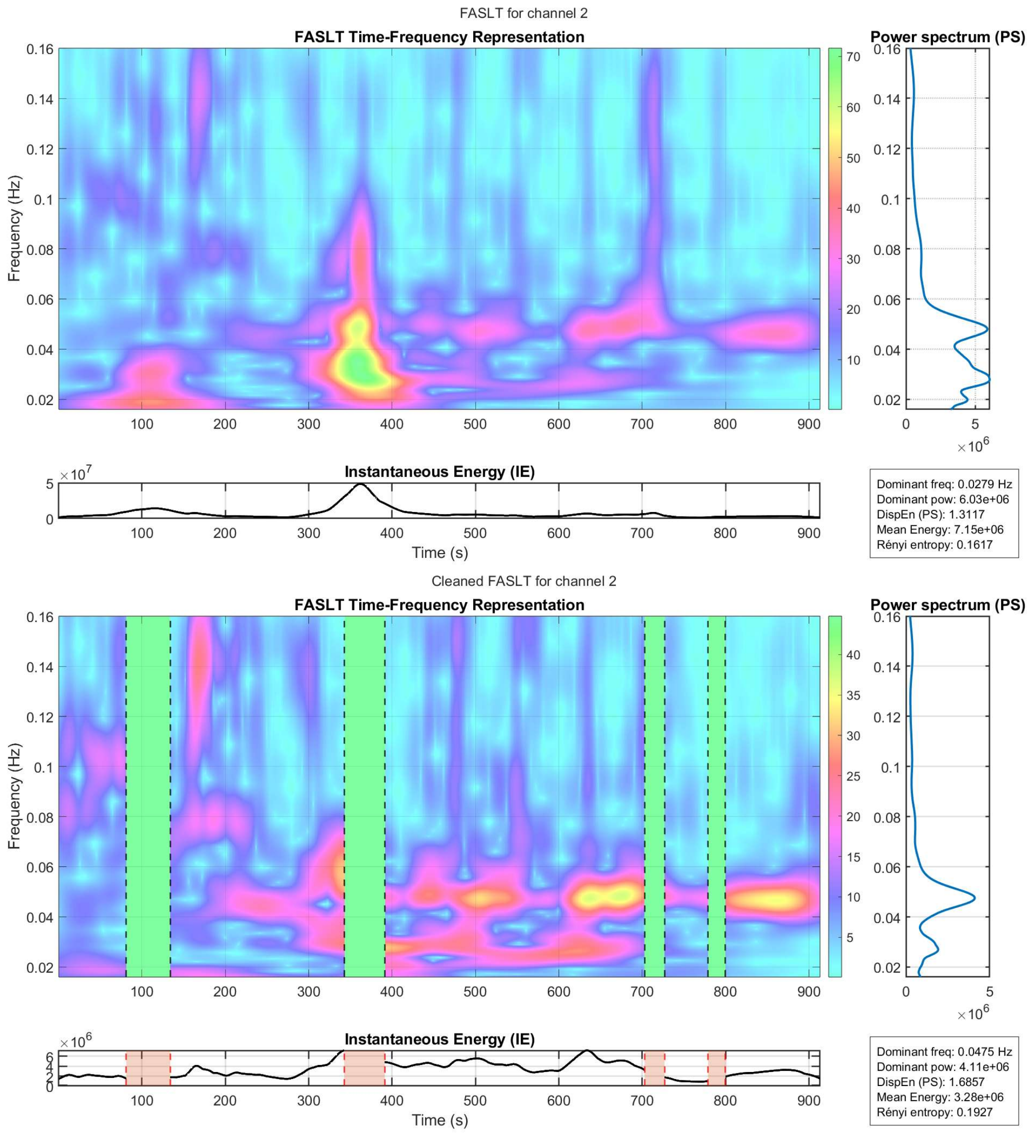


*Figure S6a: TOP:  A real EGG recording with multiple movements artifacts or electronic spikes, like the one at 100 seconds or the powerful one at around 350 seconds. The dominant frequency for this TFR is outside normogastria.. BOTTOM after the artifact removal algorithm, the signal generating Figure S8a is now represented by this new TFR. A clear normogastric dominant frequency emerges.*

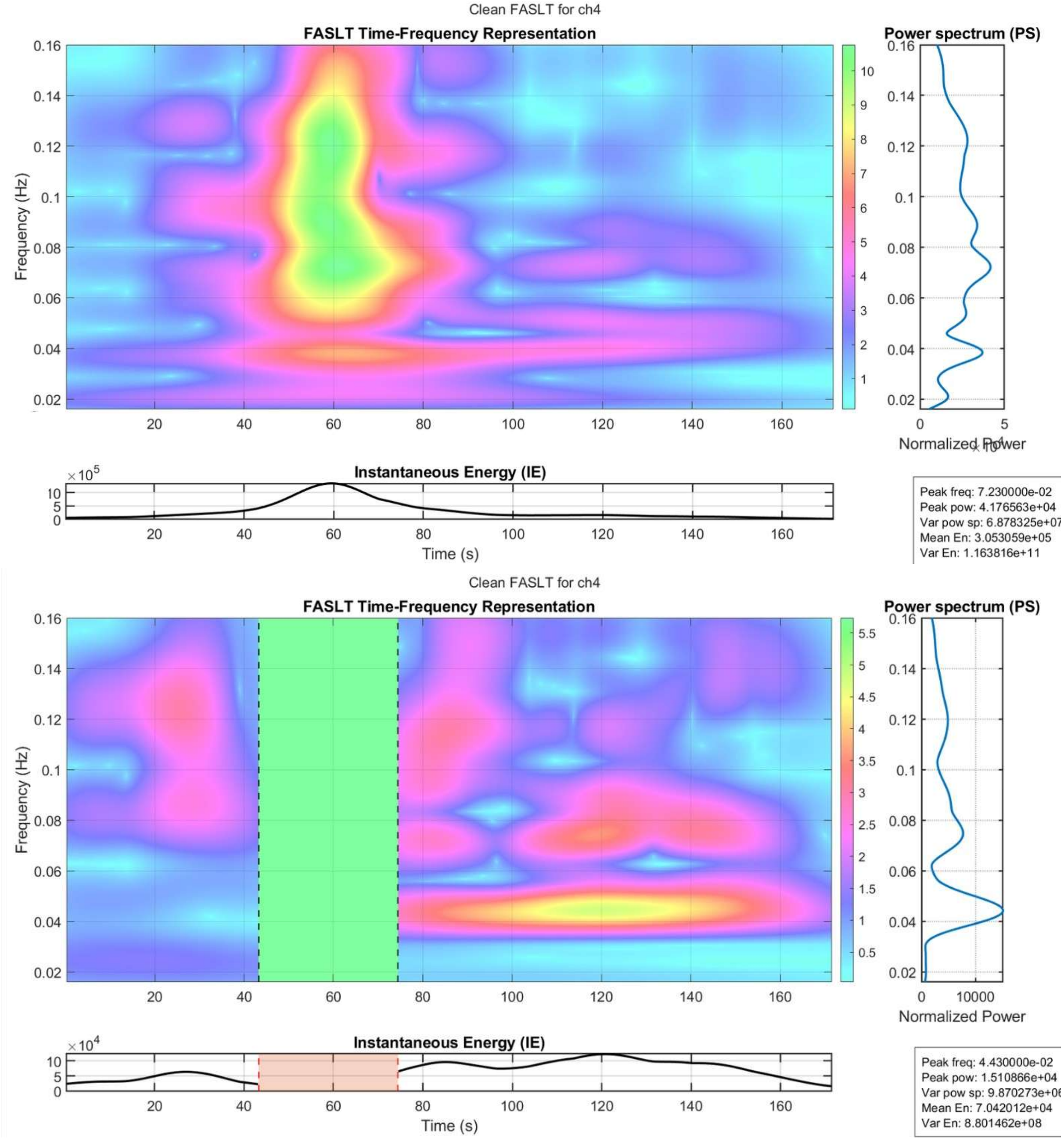


*Figure S6b: TOP: A real EGG recording a clear artifact at around 60 seconds. The dominant frequency for this TFR is outside normogastria. BOTTOM: after the artifact removal algorithm, the signal generating Figure S6b is now represented by this new TFR. A clear normogastric dominant frequency emerges.*

#### C. EFFICACY OF THE ARTIFACT REJECTION ALGORITHM: SNR

Figure S7 shows the raincloud plots comparing processing pipelines on the peak-to-background power ratio (see Main Text, Results, Section C).

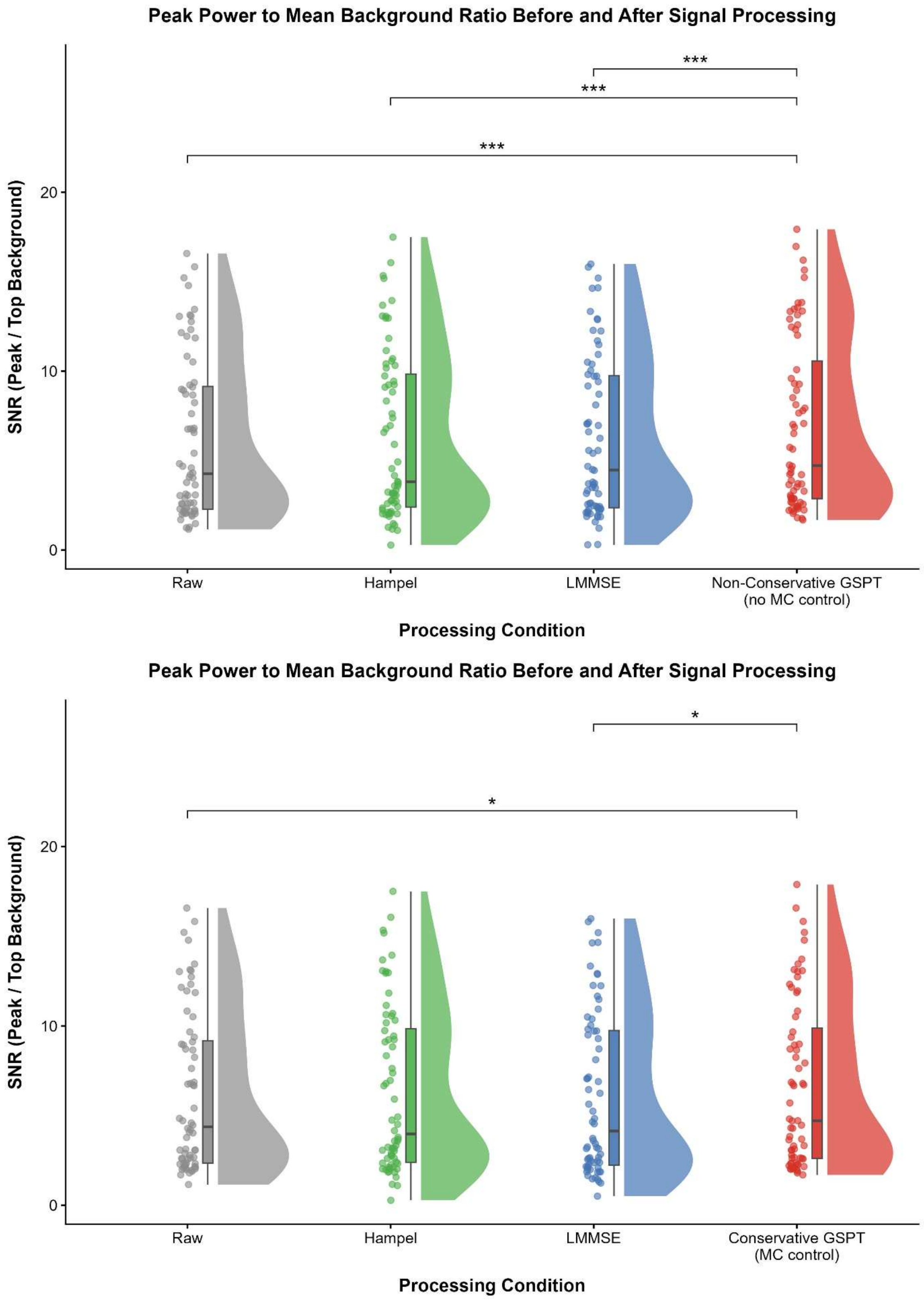


*Figure S7: Comparison of the peak-to-mean background power ratio across raw, LMMSE-processed, and different versions of GSPT-processed data containing controlled movement artifacts. (TOP): GSPT without preliminary MC control. (BOTTOM): GSPT with MC control.*

#### D. FINGERPRINT CORRELATIONS

The results on single-channel fingerprint correlations (see Main Text, Results, Section D) are plotted as violin plots in Figure S8.

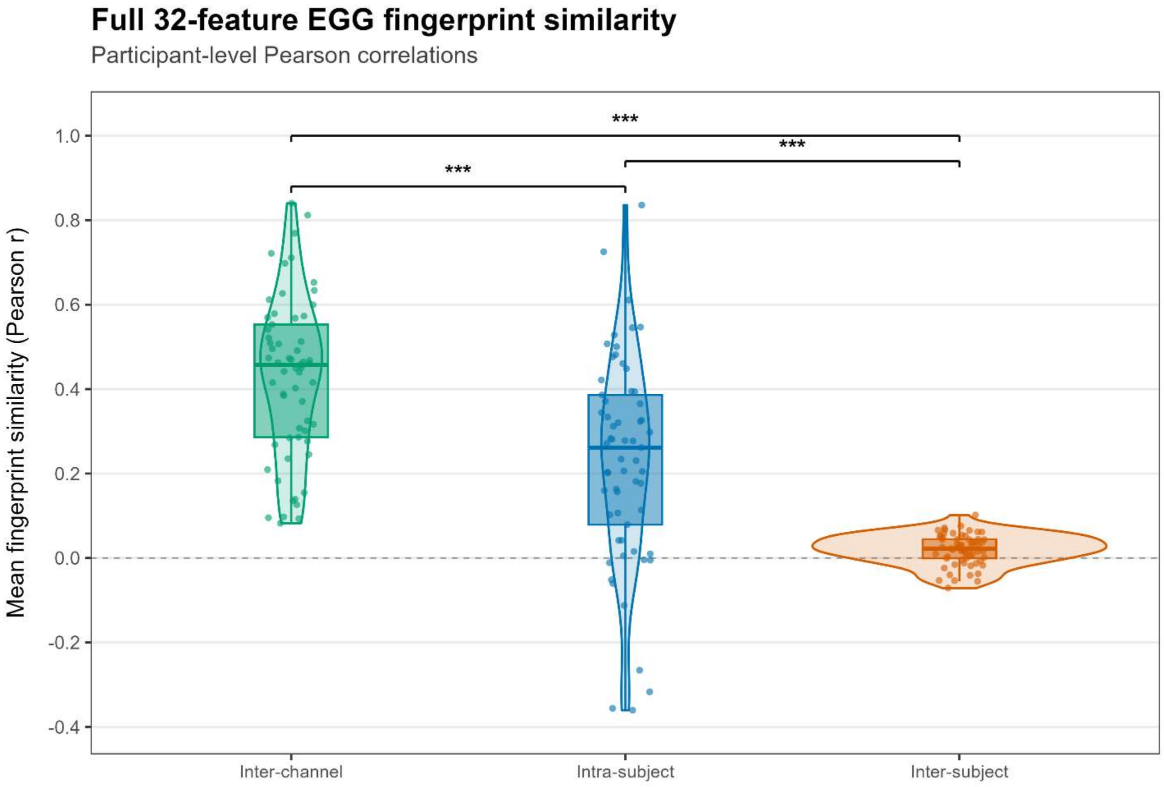


*Figure S8: Violin and box plots showing intra-subject, inter-subject, and inter-channel correlations. Brackets indicate the three planned participant-block bootstrap contrasts; *** Holm-adjusted p < 0.001. Higher Pearson correlations indicate greater fingerprint similarity. Inter-channel correlation is significantly higher than intra-subject correlation, which in turn is significantly higher than inter-subject correlation.*

#### E. QUALITY CHECK

Figure S9 shows the raincloud plots of the signal-to-noise ratio (SNR) values, comparing signals included versus excluded by the MC quality check (see Main Text, Results, Section E).

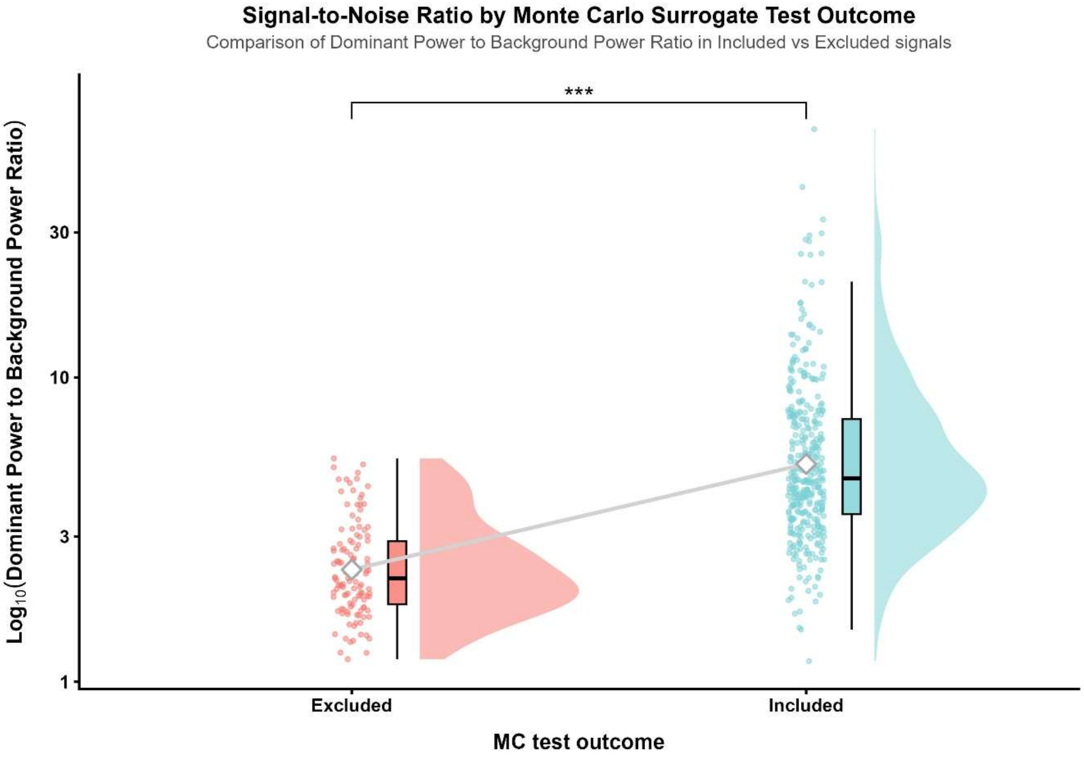


*Figure S9: Raincloud plot of the ratio between the power around the dominant frequency (± 0.015 Hz) and the rest of the gastric frequency band, comparing signals that failed the MC test (Excluded) and those that successfully passed it (Included).*

Figure S10 shows the effect of DF consistency test, by evaluating the percentage of DF in each gastric band.

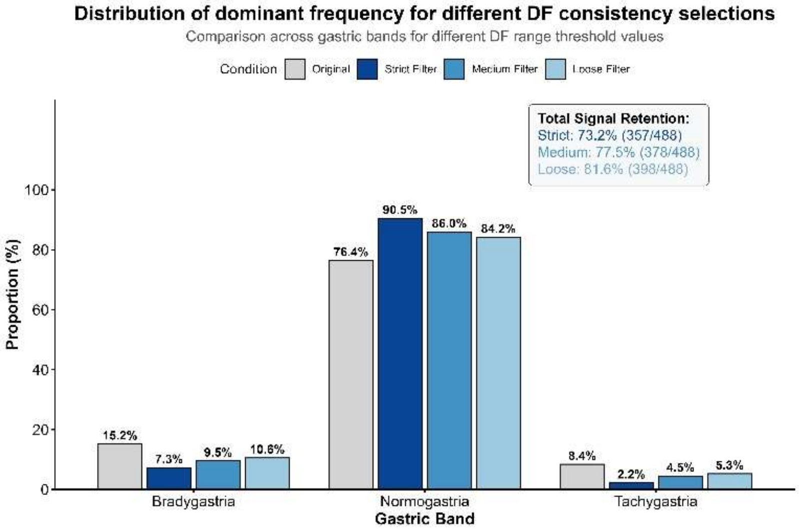


*Figure 10: Proportional distribution of dominant frequencies across traditional gastric bands (bradygastria, normogastria, tachygastria) in the original dataset compared to datasets filtered using strict, medium, and loose thresholds.*

Figure S11 (from the same section of the main text) illustrates the retention rates for the DF consistency method across three tolerance thresholds—0.005 Hz (strict), 0.010 Hz (medium), and 0.020 Hz (loose)—for each standard gastric band.

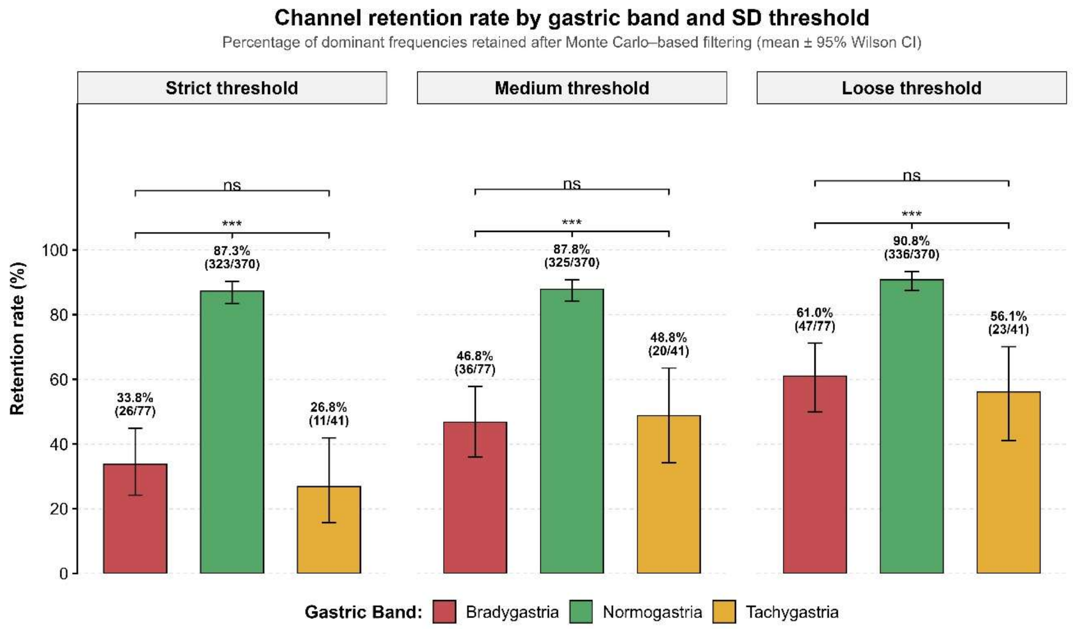


*Figure S11: Impact of DF consistency on the distribution and retention of gastric dominant frequencies. Channel retention rates by gastric band within each threshold condition. Error bars represent 95% Wilson confidence intervals.*

#### F. NON NORMOGASTRIC DF AFTER RECORDING SELECTION

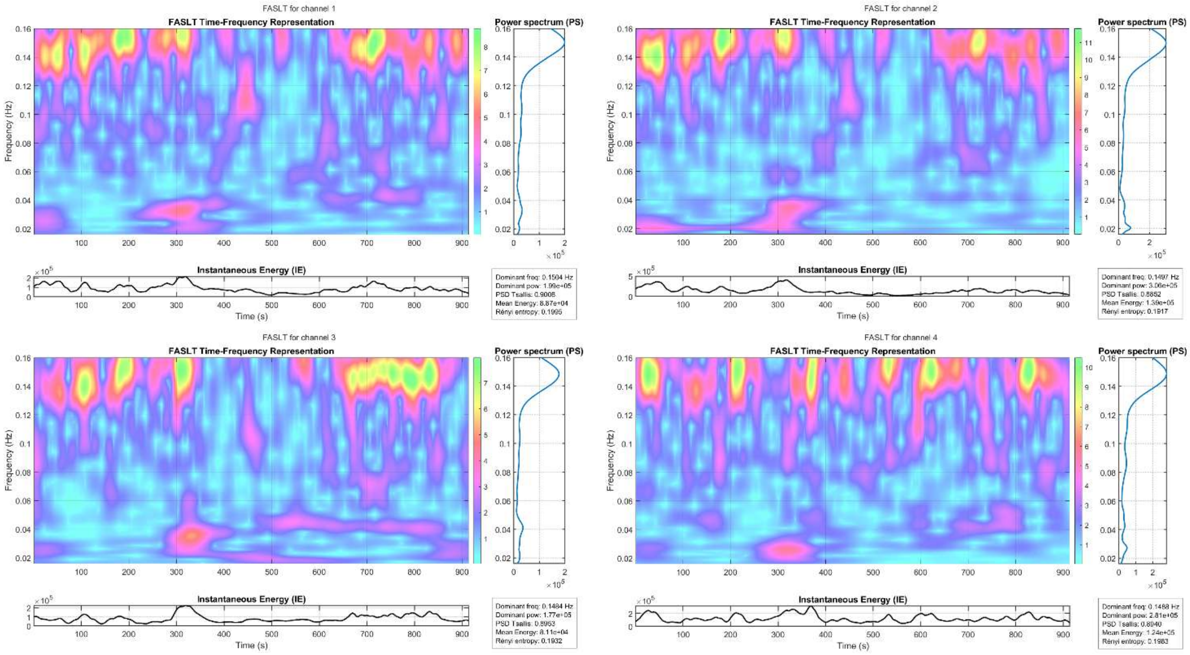


*FIgure S12: Time-frequency representations (TFRs), instantaneous energy, and power spectral densities (PSDs) for channels 1 to 4 of subject 39 (arranged clockwise from top-left). A coherent tachygastric dominant frequency (DF) emerges across the PSDs of all channels.*

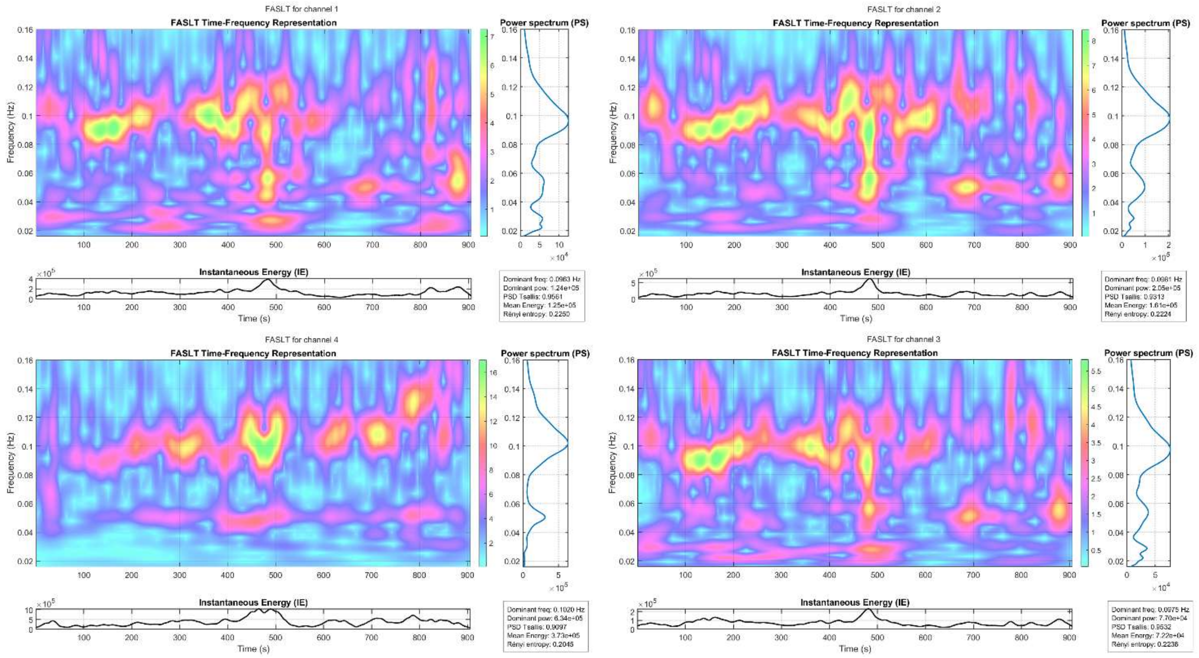


*FIgure S13: Time-frequency representations (TFRs), instantaneous energy, and power spectral densities (PSDs) for channels 1 to 4 of subject 51 (arranged clockwise from top-left). A coherent tachygastric dominant frequency (DF) emerges across the PSDs of all channels.*

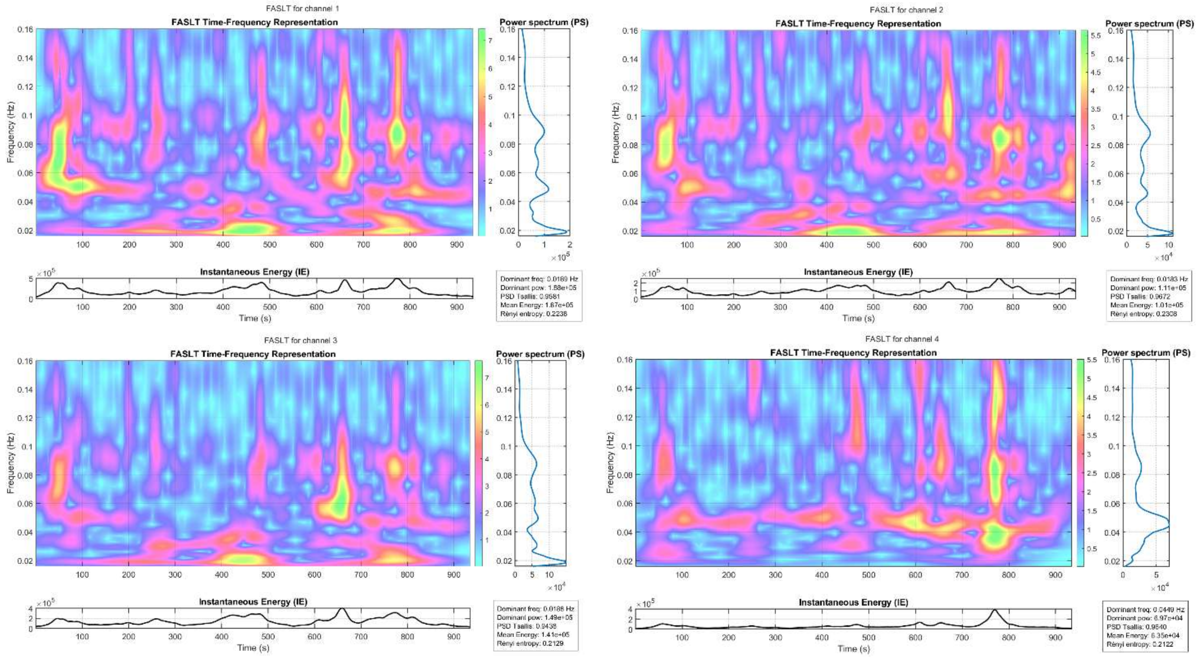


*Figure S14: Time-frequency representations (TFRs), instantaneous energy, and power spectral densities (PSDs) for channels 1 to 4 of subject 55 (arranged clockwise from top-left). A coherent bradygastric dominant frequency (DF) emerges across the PSDs of three out of four channels.*

### IX. GSPT OUTPUTS: INSTANTANOEUS CURVES AND TOTAL CORRELATIONS

This section presents examples of the GSPT outputs. Figure S11 illustrates the instantaneous amplitude, frequency, and phase derived from a single channel of a participant's clean EGG recording, whereas Figure S12 displays a spider plot of the total correlations and multivariate entropies obtained using all four recording channels.

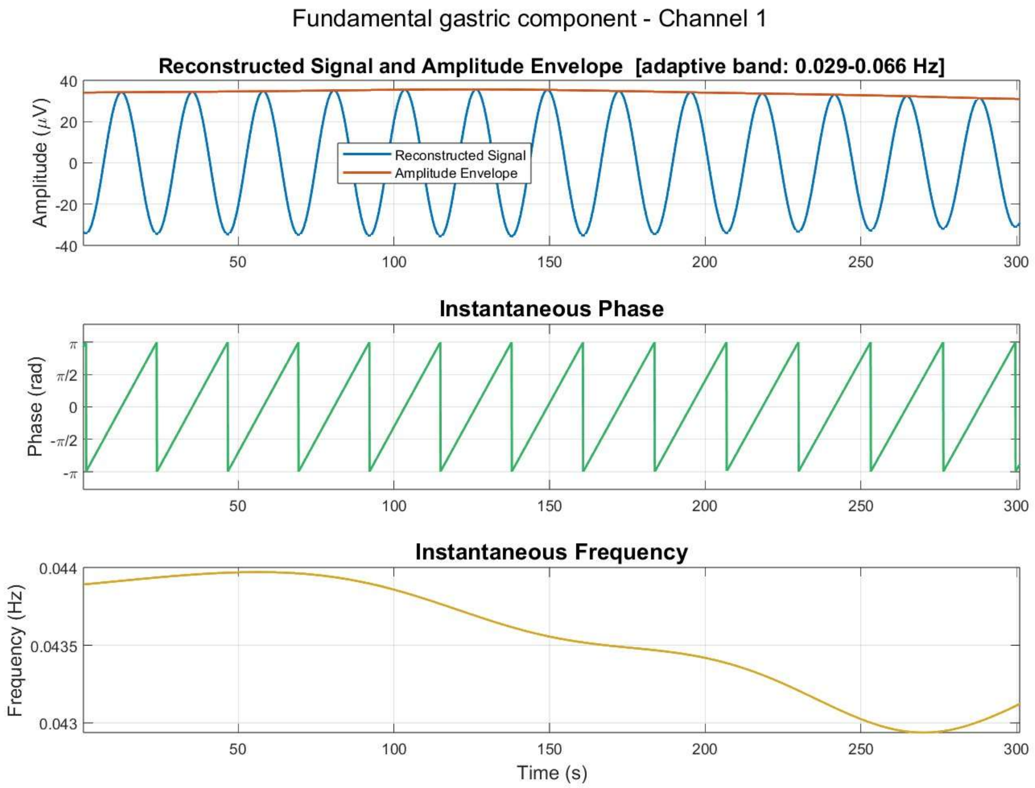


*Figure S15: Reconstructed fundamental gastric component from a real EGG recorded signal together with its 3 instantaneous quantities: instantaneous amplitude, phase and frequency. The reconstructed signal (blue line) is obtained using the formula* $s(t) = A(t) \cdot cos(2\pi\phi(t)))$*, where A(t) represents the instantaneous amplitude (red curve) and* $\phi$*(t) represents the instantaneous phase (yellow curve).*

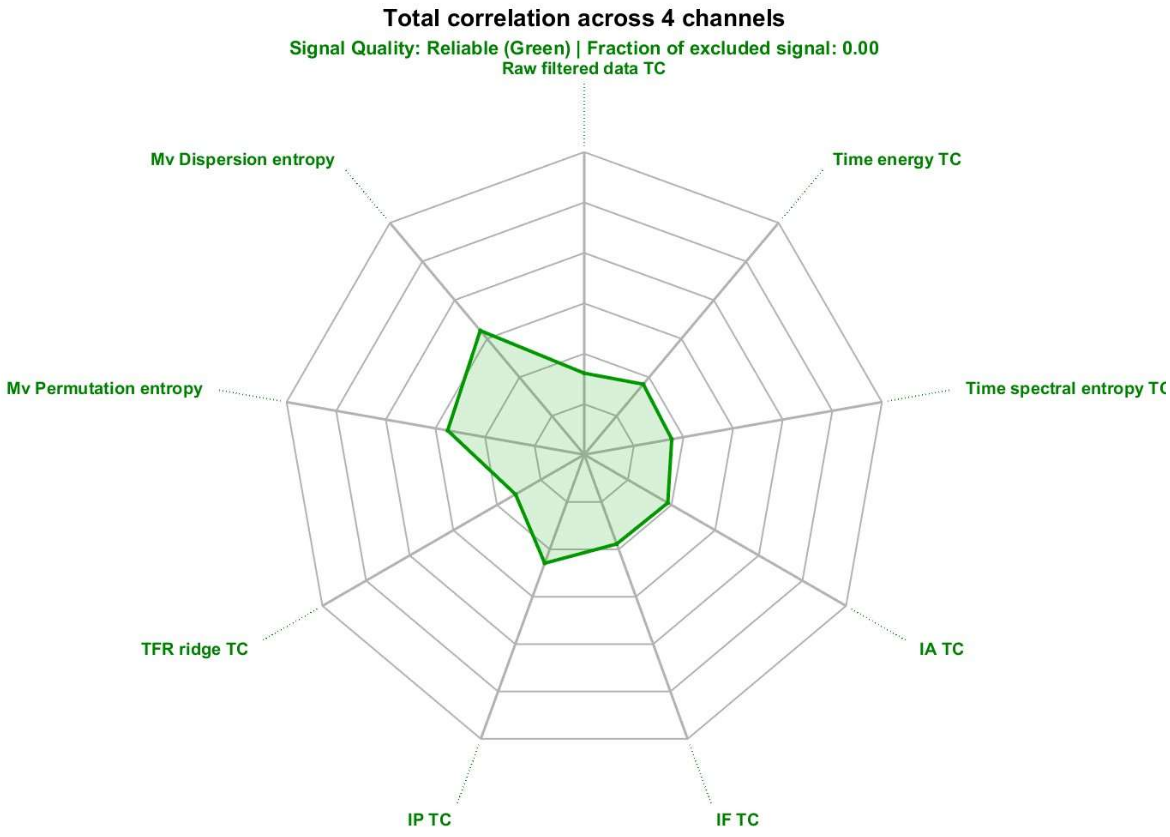


Figure S*16*: Spider plot displaying the seven total correlations and two multivariate entropies computed by the GSPT across all four channels, for the same participant presented in Figure S11.

### BIBLIOGRAPHY

[1] Tsallis, C. (1988). Possible generalization of Boltzmann-Gibbs statistics. *Journal of statistical physics*, *52*(1), 479-487.

[2] The MathWorks Inc. (2026). *Spectral Descriptors*. MATLAB & Simulink. https://it.mathworks.com/help/audio/ug/spectral-descriptors.html

[3] Flood, M. W., & Grimm, B. (2021). EntropyHub: An open-source toolkit for entropic time series analysis. PloS one, 16(11), e0259448.

[4] Yu, S., Giraldo, L. G. S., Jenssen, R., & Principe, J. C. (2019). Multivariate Extension of Matrix-Based Rényi' α-Order Entropy Functional. IEEE transactions on pattern analysis and machine intelligence, 42(11), 2960-2966.

[5] Jurdana, V. (2024). Local Rényi entropy-based Gini index for measuring and optimizing sparse time-frequency distributions. Digital signal processing, 147, 104401.

[6] Jiang, R., & Shang, P. (2025). Gradient entropy (GradEn): The two dimensional version of slope entropy for image analysis. arXiv preprint arXiv:2502.18516.

[7] Weber, I., & Oehrn, C. R. (2022). NOLITIA: an open-source toolbox for non-linear time series analysis. Frontiers in neuroinformatics, 16, 876012.

[8] Hurley, N., & Rickard, S. (2009). Comparing measures of sparsity. *IEEE Transactions on Information Theory*, *55*(10), 4723-4741.
